# Short periods of decreased water flow may modulate long-term ocean acidification in reef-building corals

**DOI:** 10.1101/2024.02.23.581783

**Authors:** Catarina P.P. Martins, Susana M. Simancas-Giraldo, Patrick Schubert, Marlene Wall, Christian Wild, Thomas Wilke, Maren Ziegler

## Abstract

Ocean acidification (OA) poses a major threat to reef-building corals. Although water flow variability is common in coral reefs and modulates coral physiology, the interactive effects of flow and OA on corals remain poorly understood. Therefore, we performed a three-month OA experiment investigating the effect of changes in flow on coral physiology. We exposed the reef-building corals *Acropora cytherea*, *Pocillopora verrucosa*, and *Porites cylindrica* to control (pH 8.0) and OA (pH 7.8) conditions at moderate flow (6 cm s^-1^) and monitored OA effects on growth. Throughout the experiment, we intermittently exposed all corals to low flow (2 cm s^-1^) for 1.5 h and measured their photosynthesis:photosynthesis (P:R) ratio under low and moderate flow. On average, corals under OA calcified 18 % less and grew 23 % less in surface area than those at ambient pH. We observed species-specific interactive effects of OA and flow on coral physiology. P:R ratios decreased after 12 weeks of OA in *A. cytherea* (22 %) and *P. cylindrica* (28 %) under moderate flow, but were unaffected by OA under low flow. P:R ratios were stable in *P. verrucosa*. These results suggest that short periods of decreased water flow may modulate OA effects on some coral species, indicating that flow variability is a factor to consider when assessing long-term effects of climate change.

## 1. INTRODUCTION

Ocean acidification (OA) constitutes a main future threat to reef-building corals (Doney et al. 2009), typically reducing their calcification rate (Kroeker et al. 2013). However, the effect of OA on respiration and photosynthesis of reef-building corals, i.e., key processes of coral physiology, may vary depending on environmental conditions and species. Although meta-analyses show that net photosynthesis generally remains stable under OA conditions (Kroeker et al. 2013; Godefroid et al. 2022), it has also been reported to decrease (e.g., Reynaud et al. 2003; Bedwell-Ivers et al. 2017) or increase (e.g., Comeau et al. 2018; Biscéré et al. 2019). Coral reefs are highly dynamic ecosystems where local environmental variability modulates coral physiology (McLachlan et al. 2021). For instance, natural diel oscillation of seawater pH in reefs (Hannan et al. 2020) may cause differences in the physiological response of coral species to OA and further complicate the assessment of its effects (Comeau et al. 2014a; Bedwell-Ivers et al. 2017; Enochs et al. 2018).

Additional physical factors such as water flow, which vary on short time scales, may underlie the complex responses of corals to OA. Water flow in coral reefs–typically high-energy ecosystems exposed to currents, waves, and tides (Sheppard et al. 2018)–can vary greatly within and among reefs (Lowe and Falter 2015). Flow velocities vary even within single reef locations (e.g., 0–30 cm s^-1^; Roik et al. 2016), with temporal decreases during the diel cycle associated with tides (Green et al. 2018; Lindhart et al. 2021). Flow also differs between reef environments, with back-reef environments typically experiencing lower flow than reef-crest environments (Madin et al. 2006). Periods of low flow may be relatively common in some reefs (e.g., Hench et al. 2008), and flow immediately adjacent to coral colonies is further reduced due to the formation of recirculation zones in complex reef topographies (Hench and Rosman 2013).

In reef-building corals, water flow modulates respiration and photosynthesis (Dennison and Barnes 1988; Patterson et al. 1991) with species-specific effects (e.g., Rex et al. 1995; Schutter et al. 2010). Differences in physiological responses between species to water flow may be associated with its effect on the coral boundary layer. This is the layer of seawater bordering the coral surface that controls mass transfer between the coral and bulk seawater (Atkinson and Bilger 1992; Shashar et al. 1993). Water flow variability also has implications for coral physiology. For instance, photochemical efficiency may be higher under alternating high-to-low flow conditions than under constant flow (Smith and Birkeland 2007). Thus, speed and short-term variability of water flow elicit complex patterns in coral physiology.

Although water flow is a prevailing characteristic of all coral reefs, knowledge of the interactive effects of different flow regimes and OA on reef-building corals remains limited (Noisette et al. 2022). While high-flow environments have been proposed as refuges from the effects of ocean warming (Fifer et al. 2021), low-flow environments are currently considered potential refuges from OA for calcifying organisms (Hurd 2015). Effects of flow and OA on coral physiology may be complex, potentially interactive, and differ by exposure time. For instance, during 1-h short-term OA exposure, net photosynthesis was similar between low (1 cm s^-1^) and moderate water flow (4–13 cm s^-1^; Osinga et al. 2017). Similarly, after two-day exposure to OA, coral communities briefly exposed to high (35 cm s^-1^) and moderate flow (8 cm s^-1^) also had similar net photosynthesis under OA (Anthony et al. 2013). Whereas, after two-month exposure to OA and different flow regimes (2.5 or 8 cm s^-1^), net photosynthesis under low flow was decreased in *Acropora yongei* and increased in *Plesiastrea versipora* (Comeau et al. 2019a). Similarly, decreased calcification of reef-building corals under OA may be alleviated by temporary 24-h exposure to moderate water flow (5 and 10 cm s^-1^) compared to low flow (2 cm s^-1^; Comeau et al. 2014c). These results suggest that acclimatisation to OA and flow regimes may occur on different time scales. Thus, a systematic investigation of long-term OA effects on coral physiology combined with short-term fluctuations of water flow, as occur in coral reefs, is needed.

The overarching aim of this study was to assess the physiological response of three reef-building coral species, *Acropora cytherea* (Dana, 1846)*, Pocillopora verrucosa* (Ellis & Solander, 1786), and *Porites cylindrica* Dana, 1846, to changes in water flow under control and OA conditions. Specifically, we tested how OA conditions (pH 7.8) maintained for three months at constant moderate flow (6 cm s^-1^) affected I) coral calcification and surface growth compared to control conditions (pH 8.0) and II) coral photosynthesis:respiration (P:R) ratios under short periods of low flow (2 cm s^-1^) compared to responses under moderate flow. This study will help to disentangle the complex effects of OA on coral physiology and better understand the role of hydrodynamics in the response of reef-building corals to OA.

## 2. MATERIALS AND METHODS

### 2.1. Study species and experimental design

A three-month experiment was conducted to investigate the physiological response of three scleractinian coral species, *Acropora cytherea*, *Pocillopora verrucosa*, and *Porites cylindrica,* to OA together with the effect of changes in water flow conditions using respirometry assays. Coral colonies (Table S1) were maintained at the ‘Ocean2100’ long-term coral experimental facility (8,000 L closed recirculating system of artificial seawater, Table S2) at Justus Liebig University Giessen, Germany, for at least six months before the experiment. Conditions in the long-term culturing tanks (256 L) were 11:13 h light:dark photoperiod with a light intensity of 230 µmol m^-2^ s^-1^, temperature of 26.0 ± 0.5 °C, and daily feeding of a mix of frozen *Artemia* sp., *Mysis* sp., and copepods. For the experiment, a total of four colonies per species were used and cut into eight fragments using a small angle grinder (Dremel Multitool 3000-15, The Netherlands). Fragments were attached to tiles with two-component glue (CoraFix SuperFast, Grotech, Germany) and transferred to the experimental setup. Corals were acclimated to the experimental setup for five weeks before the start of the experiment. During the first 10 days of the acclimation period, corals were administered the long-term culturing feed, after which the culturing feed was provided with lower frequency (2.7 mg L^-1^ of frozen copepods every two days) due to the lower density of coral fragments within experimental tanks. Corals also received constant dissolved nutrient conditions via the connected water system.

Experimental treatments consisted of two pH levels, with the control treatment mimicking present-day atmospheric *p*CO2 concentration on some reefs (∼500 µatm *p*CO2; Ziegler et al. 2021), and the OA treatment with values projected in the long term (2081–2100) for surface ocean pH in coral reefs (UNEP-WCMC et al. 2021) under SSP2-4.5 (0.20 pH units lower relative to 1961–1990; Iturbide et al. 2022; IPCC 2023). The physiological response of the corals was monitored throughout both the acclimation and experimental period, which were conducted from 15 October 2019 to 28 February 2020. Buoyant weight was measured three weeks before the start of gradual pH decrease (t-3) and after 13 weeks under OA (t13), while surface area was documented four weeks before the start of gradual pH decrease (t-4) and after 14 weeks under OA (t14). The time points of these two parameters differed by one week due to the duration of measurements (buoyant weight measurement of all coral fragments took three days and surface area six days). P:R ratios were measured in low (2 cm s^-1^) and moderate flow (6 cm s^-1^) four times on all fragments (32 fragments per species per flow condition = 192 fragments per time point). Measurements of all fragments took six days to complete per time point and were performed once at the end of the acclimation period (t-1, one week before the start of gradual pH decrease) and three times during the experimental period (t3, after three weeks under OA; t7, after seven weeks; t12, after 12 weeks). A complete timeline of all physiological measurements can be found in Fig. 1A and details of the physiological assessments are outlined below.

**Fig. 1.**
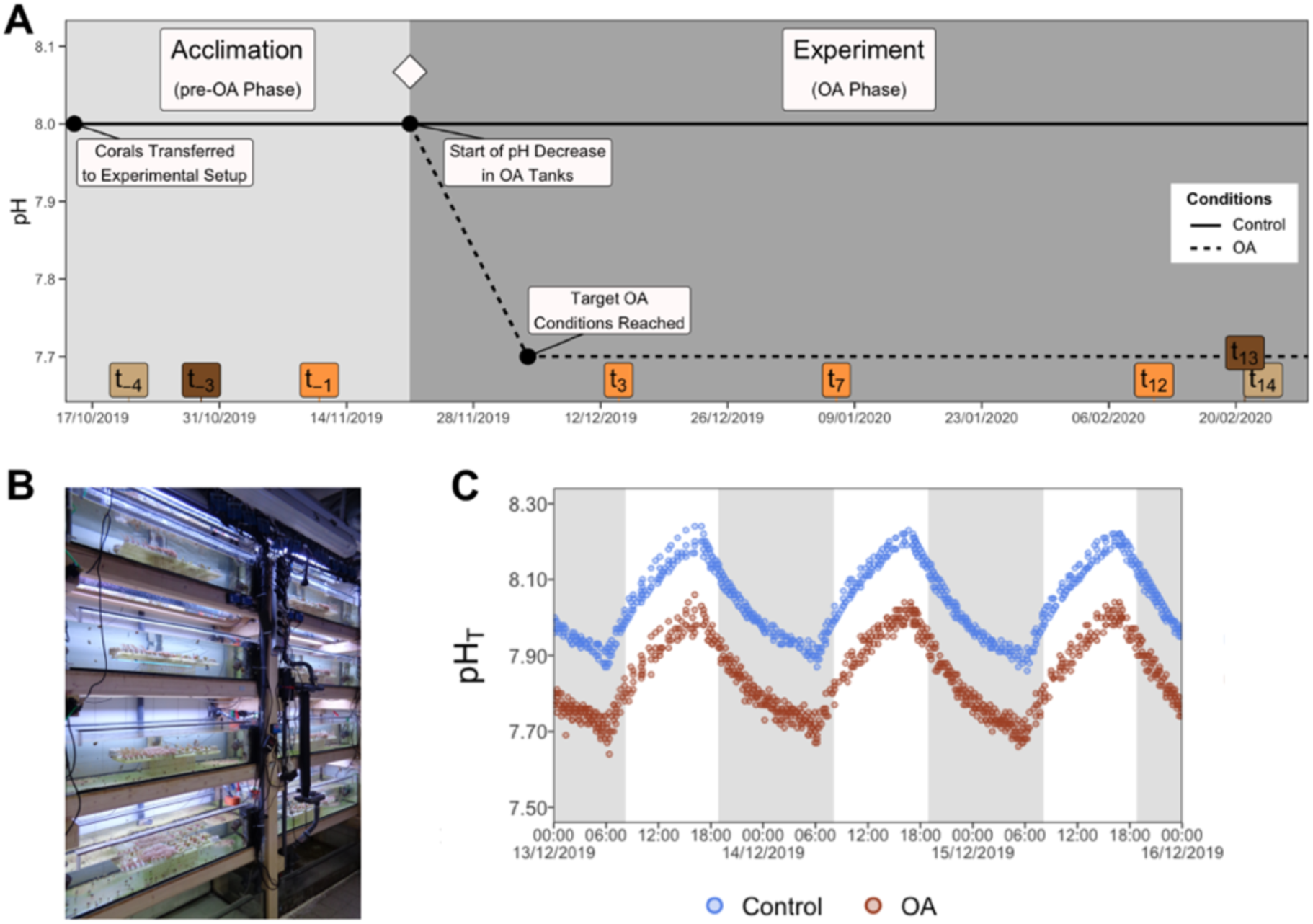
Experimental design and setup. (A) Timeline of physiological measurements performed throughout the acclimation and experimental periods. Different colours indicate different types of physiological measurements (calcification, dark brown; surface growth, light brown; photosynthesis:respiration ratio, orange). The white diamond indicates the start of the experiment. Dates are provided as DD/MM/YYYY. (B) Overview of the experimental setup. See Fig. S1 for a detailed close-up of an individual experimental tank. (C) Snapshot of diel pH oscillation during the experiment. Shaded grey areas indicate nighttime. OA, ocean acidification; t_-1_, one week before the start of gradual pH decrease; t_-3_, three weeks before; t_-4_, four weeks before; t_3_, after three weeks under OA conditions, including two weeks of gradual pH decrease; t_7_, after seven weeks; t_12_, after 12 weeks; t_13_, after 13 weeks; t_14_, after 14 weeks; pH_T_, pH on the total scale

### 2.2. Experimental setup and treatment conditions

The experimental setup consisted of eight 120 L tanks divided into two experimental pH treatments (four tanks per treatment, 16 fragments per species per treatment) (Fig. 1B). Each tank housed one fragment per colony (total of four fragments per species in each tank) with 15 cm spacing between them in the direction of flow. In addition, experimental tanks contained other scleractinian and octocorals with the same number of individuals per tank (Fig. S1A). The experimental tanks were supplied with water from the 8,000 L closed recirculating system of artificial seawater (calcium concentration: 396 ± 6 mg L^-1^, phosphate: < 0.02 mg L^-1^, nitrate: < 0.02 mg L^-1^, nitrite: < 0.01 mg L^-1^) with an inflow rate of 20–40 L h^-1^ (corresponding to a 100 % tank volume turnover every 3–6 h). In addition, the large water system received weekly water changes of ∼10 % of the water volume. Temperature was maintained at 26 °C through a feedback-controlled heater (300 W; 548, Schego, Germany). Water flow conditions, consisting of a flow velocity of 6 cm s^-1^ (measured at the position of coral fragments; OTT MF pro, OTT Hydromet GmbH, Germany) and a standing wave with an amplitude of 5 mm, were generated with two circulating pumps (ES-28, Aqualight, Germany) and one wave generator (6208, Tunze, Germany). Salinity in the tanks was monitored daily using a conductivity sensor (TetraCon 925, WTW, Germany) and maintained at 35. Light was provided by two T5 bulbs (54 W, Aqua-Science, Germany), producing a light intensity of 176 ± 31 µmol m^-2^ s^-1^ in an 11:13 h light:dark photoperiod. Light intensity in the experimental tanks differed slightly from conditions in the culturing tanks due to technical reasons.

Seawater pH was constantly monitored using a digital controller (Profilux 3, GHL, Germany) attached to pH electrodes in each tank (GHL, Germany), which were calibrated every two weeks using NBS buffers. Values of pHNBS were converted to total scale (pHT) using equations from Millero (2013) and Takahashi et al. (1982). pH values are expressed in total scale throughout the text. OA conditions in each treatment tank were generated individually via pH-controlled CO2 dosing (bubbling) using solenoid valves, which controlled the release of CO2. Pumping was done through one of the circulating pumps to aid CO2 dissolution and dispersion in seawater. pH was gradually decreased in OA treatment tanks and lowered by 0.01–0.02 units every day over two weeks until target values were reached (Fig. S2A). Target OA conditions were maintained for 86 days, including diel oscillation of pH mimicking naturally occurring variability (Hannan et al. 2020; Ziegler et al. 2021). Total alkalinity (TA) was measured in each experimental tank by open-cell potentiometric titration using a titrator (TitroLine 7000, SI Analytics, Germany) equipped with a glass pH-combination electrode (A 162 2M-DIN-ID, SI Analytics, Germany). Measurements were made following SOP3b (Dickson et al. 2007) on 50 g samples with 0.1N HCl (Titrisol, Merk, Germany) in 35 g L^-1^ NaCl and corrected using certified reference materials (Batch 183, A.G. Dickson Laboratory, Scripps Institution of Oceanography, UCSD, USA; Dickson et al. 2003). Measurements were performed every 2–4 days during the first two weeks of the experiment and then every 1–2 weeks. TA was calculated using a modified Gran approach (Millero 2013). Alkalinity was also monitored daily and maintained with two automatic in-house constructed calcium reactors (pH 6.2–6.4, coral rubble) and dosing of NaHCO3 in a common reservoir tank. The calcium reactor was feedback controlled by an alkalinity controller (Alkatronic, Focustronic, Hong Kong) based on 3-hourly automatic titrations. Seawater carbonate chemistry was calculated from days with TA measurements using pHNBS and temperature values for a whole day and the corresponding value of TA and salinity. TA and salinity values were assumed to be representative of the conditions of the entire day on which they were measured. Calculations were performed in the program CO2SYS (v25; Pelletier et al. 2007) with carbonic acid dissociation constants from Mehrbach et al. (1973) refit by Dickson and Millero (1987). This approach is suitable for biological OA experiments with treatments that have differences larger than 100 µatm *p*CO2 (Watson et al. 2017) and allowed us to account for the diel oscillation of pH in the tanks.

### 2.3. Coral calcification and surface growth determination

Calcification was determined from measurements of buoyant weight (Jokiel et al. 1978) and calculated as the difference between final (t13) and initial (t-3) buoyant weight (accuracy 0.001 g; KB 360-3N, KERN & Sohn GmbH, Germany), converted to dry weight using an aragonite density of 2.93 g cm^-3^ (Spencer Davies 1989), and normalised by initial surface area (t-4) and time. Surface growth was determined using 3D scanning (Artec Spider 3D with Artec Studio 9, Artec 3D, Luxembourg) by documenting coral surface area, following Reichert et al. (2016). Briefly, corals were placed on a rotating plate and scans were captured in air within 60–90 seconds. 3D models were calculated by performing fine serial registration, global registration (minimal distance: 10, iterations: 2,000, based on texture and geometry), sharp fusion (resolution: 0.2, fill holes by radius, max. hole radius: 5), and outlier removal (*A. cytherea* and *P. cylindrica*, SD: 3; *P. verrucosa*, SD: 4). Artefact objects were removed (small objects filter). To determine coral surface areas, meshes were trimmed manually at the tissue border. All meshes were exported as Wavefront “.obj” files to MeshLab Visual Computing Lab-ISTI-CNR (v1.3.4, BETA, 2014; Cignoni et al. 2008), and surface area was calculated using the “compute geometric measures” tool. Surface growth rates were determined as the difference between final (t14) and initial (t-4) surface area and normalised by initial surface area and time.

### 2.4. Measurement of photosynthesis:respiration ratios

P:R ratios were derived from oxygen production and consumption rates at low and moderate flow conditions measured at the same time of day to avoid bias due to diurnal variation. Coral fragments were incubated individually in sealed 1 L glass chambers for 90 min at 191 ± 23 µmol m^-2^ s^-1^ to measure oxygen production, followed by 90 min in darkness to measure oxygen consumption. Chambers were filled with seawater from the corresponding treatment and maintained at 26 °C. Low and moderate water flow conditions in the incubation chambers were generated with a magnetic stirring bar (Fig. S1B) (Rades et al. 2022) and measured by visual tracking of small, neutrally buoyant plastic beads. Dissolved oxygen concentration was measured in each chamber at the start and end of each incubation using an optical oxygen sensor (FDO 925, WTW, Germany). Four empty chambers were included in every incubation run to control for background biological activity. Rates of oxygen production and consumption were calculated as the change in dissolved oxygen per incubation volume (calculated as the difference between water volume and volume of the coral fragment with its tile) and normalised to incubation time. P:R ratios were calculated as the ratio of gross oxygen production (i.e., the sum of net oxygen production and consumption) to oxygen consumption with an 11:24 h metabolic cycle (i.e., the ratio of total oxygen produced during daylight hours to that consumed during a 24 h period) to estimate daily autotrophic capability (McCloskey et al. 1978). Rates of respiration and photosynthesis were not analysed in this study due to the presence of inconclusive patterns between rates measured during the acclimation period (t-1) and at the end of the experimental period (t12), but are available for inspection in Fig. S3 and Table S3. Additional details are provided as Supplementary Text.

### 2.5. Statistical analysis

All statistical analyses were performed in R (v.4.1.0; R Core Team 2021) using RStudio (v1.4.1103; RStudio Team 2021). All plots were produced using the R package *ggplot2* (Wickham 2016). Changes in the physiological parameters of the three studied corals were investigated using linear mixed-effects models (LMMs). Differences between species in calcification and surface growth rates were assessed using LMMs with species (3 levels: *A. cytherea*, *P. verrucosa*, and *P. cylindrica*) as a fixed factor, and colony and treatment as random factors, while differences in P:R ratios were analysed using the same model structure but with the addition of coral fragment identity (ID) as a random effect. To test the effect of OA on calcification and surface growth, we used LMMs constructed for each species with treatment (2 levels: control and OA) as a fixed factor and colony as a random factor. The P:R ratio response to OA and flow over time was assessed using LMMs constructed for each species with treatment (2 levels: control and OA), flow (2 levels: low and moderate), and time (3 levels: t3, t7, and t12) as fixed factors in a fully crossed design, and ID, colony, and tank as random factors. LMMs were performed using the R package *lme4* (Bates et al. 2015). Model validation was performed by graphically assessing homogeneity and normality assumptions, and models were inspected for any influential observations using the R package *performance* (Lüdecke et al. 2021). The numerical output of LMMs was extracted using the R package *sjPlot* (Lüdecke 2021) and is provided with model formulas in Tables S4 and S5. We then computed type-II ANOVA tables of the fixed effects of LMMs using the Kenward-Roger approximation for the degrees of freedom in the R package *car* (Fox & Weisberg 2019. Type II sums of squares was selected to compute ANOVAs, following previously recommended protocol for assessing main effects individually in the absence of interactions (Langsrud 2003; Hector et al. 2010). Post hoc analyses were performed using the R package *emmeans* (Lenth 2021) with Bonferroni adjustment of p-values.

Differences in seawater chemistry between treatments were tested using daily mean values from days with TA measurements and the same approach as above (LMM-ANOVA) with treatment as a fixed factor (2 levels: control and OA) and tank and date as random factors.

All fragments of *A. cytherea* in one control tank bleached and subsequently died eight weeks into the experiment. The fragments of *P. verrucosa* from the same tank also showed signs of bleaching after 11 weeks into the experiment and were driving the response patterns in the data analyses (as revealed by correlation analysis, Fig. S4). Therefore, all fragments from this tank were excluded from the analyses. Based on the regular monitoring of water parameters, the underlying reason for the affected fragments could not be identified. The other fragments of these species appeared healthy. In addition, the coral fragments did not show visual signs of bleaching or necrosis in response to the OA treatment.

## 3. RESULTS

### 3.1. Ocean acidification conditions

During the three months of the experiment, pH of the control was significantly higher at 7.98 ± 0.13 (mean ± SD; daily range: 7.79–8.19) than in the OA treatment at 7.78 ± 0.13 (range: 7.60–8.01; LMM-ANOVA, F = 298, p < 0.001). The control and OA treatments had similar diel pH oscillations (Fig. 1C), with a diel range of 0.4 pH units in both treatments throughout the experiment (Fig. S2B; Table S6). *p*CO2 values were significantly lower in the control at 480 ± 171 µatm (range: 244–769 µatm) than in the OA treatment at 813 ± 286 µatm (range: 416–1,262 µatm; LMM-ANOVA, F = 273, p < 0.001). Total alkalinity and temperature were similar between treatments (LMM-ANOVA, F = 0.2/1.2, p > 0.05; Table 1). Seawater chemistry per tank and a summary of the full recording of pH and temperature values are provided in the Supplementary Material (Fig. S2B,C; Tables S6,S7).

**Table 1.**
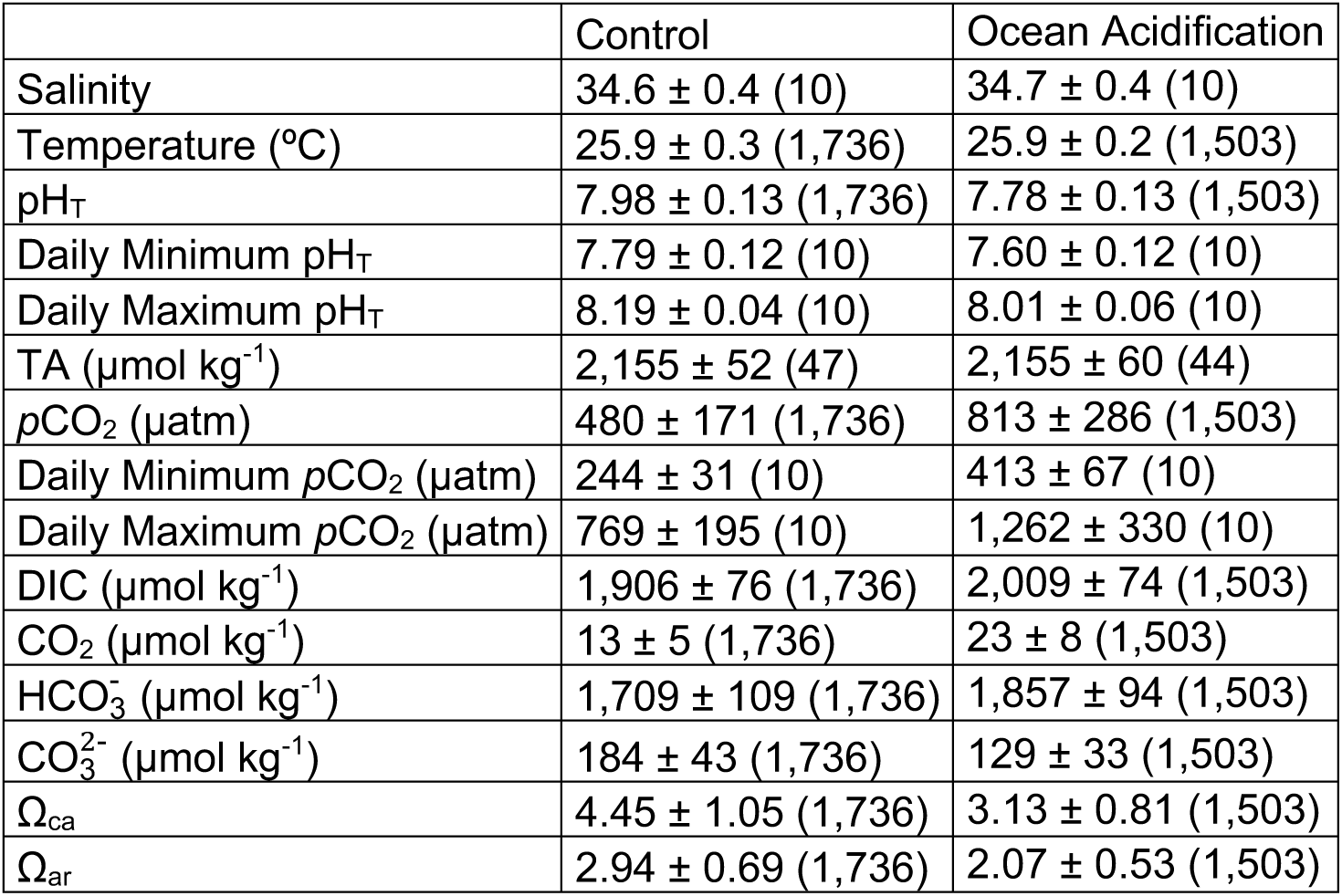
Seawater chemistry during a three-month ocean acidification experiment. Values are expressed as mean ± SD with measurement replication (n). pH_T_, pH on the total scale; TA, total alkalinity; *p*CO_2_, partial pressure of CO_2_; DIC, dissolved inorganic carbon; Ωca, calcite saturation; Ω_ar_, aragonite saturation

### 3.2. Ocean acidification decreased coral calcification and surface growth

During the experimental period, all coral fragments grew in weight and surface area, but increases differed between the three investigated species (LMM-ANOVA, F = 17.0/17.1, p < 0.01; Fig. 2). *Acropora cytherea* had a 16 % weight gain over the experiment (pooled over treatments) and a significantly lower overall calcification rate than *Pocillopora verrucosa* and *Porites cylindrica*, which presented a weight gain of 42 and 30 %, respectively (Tables S8,S9). Also, while the surface area of *A. cytherea* increased by 58 % during the experiment across treatments, it increased by 163 and 140 % in *P. verrucosa* and *P. cylindrica*, respectively, and was significantly higher than surface area growth in *A. cytherea* (Tables S8,S9).

**Fig. 2.**
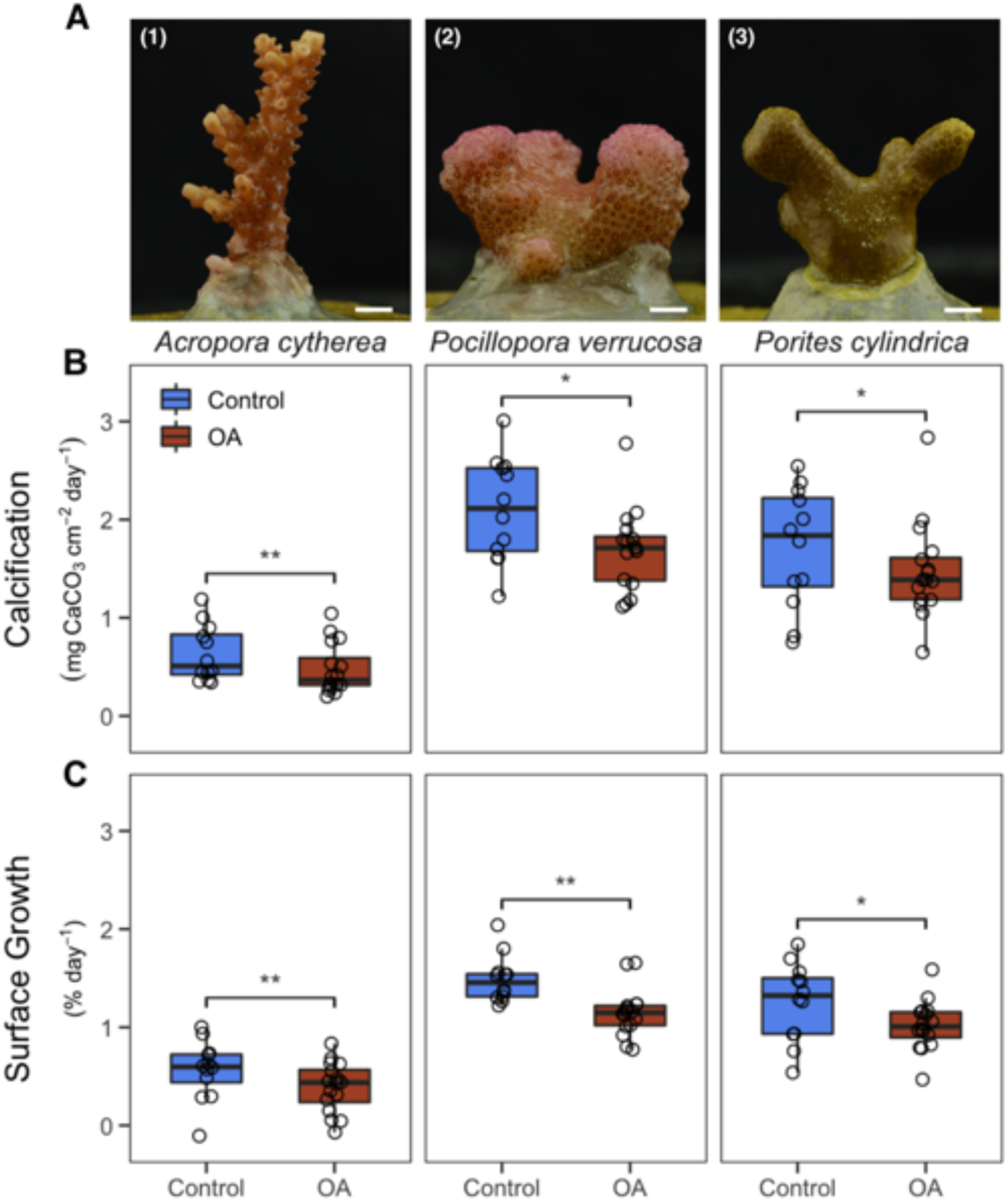
Growth effects of ocean acidification (OA) on three reef-building coral species. (A) Photographs of the investigated reef-building coral species taken at the end of the acclimation period: (1) *Acropora cytherea*, (2) *Pocillopora verrucosa*, (3) *Porites cylindrica*. Scale bar = 5 mm. (B) Calcification and (C) surface growth of *A. cytherea*, *P. verrucosa*, and *P. cylindrica* during three months in a control and OA treatment. Boxes represent the first and third quartiles with lines as medians and whiskers as the minimum and maximum values or up to the 1.5 × interquartile range (IQR), whichever is reached first. Stars indicate significant differences between the control and OA treatment (p < 0.01**, p < 0.05*, from linear mixed-effects models with ANOVA).

The OA treatment decreased calcification and surface growth rates in all species (Fig. 2B,C). *Acropora cytherea* showed a 24 and 30 % reduction in calcification and surface growth, respectively (LMM-ANOVA, F = 14.1/11.1, p < 0.01) (Table S8). In *P. verrucosa,* calcification and surface growth decreased by 20 and 23 % (LMM-ANOVA, F = 6.8/13.8, p < 0.05/0.01), and in *P. cylindrica* by 13 and 19 %, respectively (LMM-ANOVA, F = 7.3/5.8, p < 0.05) (Table S8).

### 3.3. Ocean acidification effects on photosynthesis:respiration ratios increased over time

The three investigated coral species displayed time-delayed physiological responses to OA treatments with complex interactions with water flow (Fig. 3). P:R ratios were overall similar between species during the experiment (pooled over treatments and time, LMM-ANOVA, F = 3.9, p > 0.05). They increased after the pre-OA phase (t_-1_; Fig. 3; Table S10) and differed between time points during the OA-phase for all species (pooled over treatment, LMM-ANOVA, *A. cytherea*, F = 11.1, p < 0.001; *P. verrucosa*, F = 11.3, p < 0.001; *P. cylindrica*, F = 63.0, p < 0.001; Table S11). The P:R ratio was reduced in the OA treatment compared to the control with interactive effects with time (LMM-ANOVA, Treatment-Time interaction) in *A. cytherea* (F = 13.9, p < 0.001) and *P. cylindrica* (F = 3.4, p < 0.05) under moderate flow (Fig. 3; Table S10), but not in *P. verrucosa* (F = 1.1, p > 0.05). OA effects occurred from three weeks onward in *P. cylindrica* and after 12 weeks in *A. cytherea* (Table S11). P:R ratio was on average lower under low flow than moderate flow conditions (LMM-ANOVA, *A. cytherea*, F = 6.0, p < 0.05; *P. cylindrica*, F = 20.9, p < 0.001), except in *P. verrucosa*, for which this effect was reversed (LMM-ANOVA, F = 150.9, p < 0.001) (Fig. 3).

**Fig. 3.**
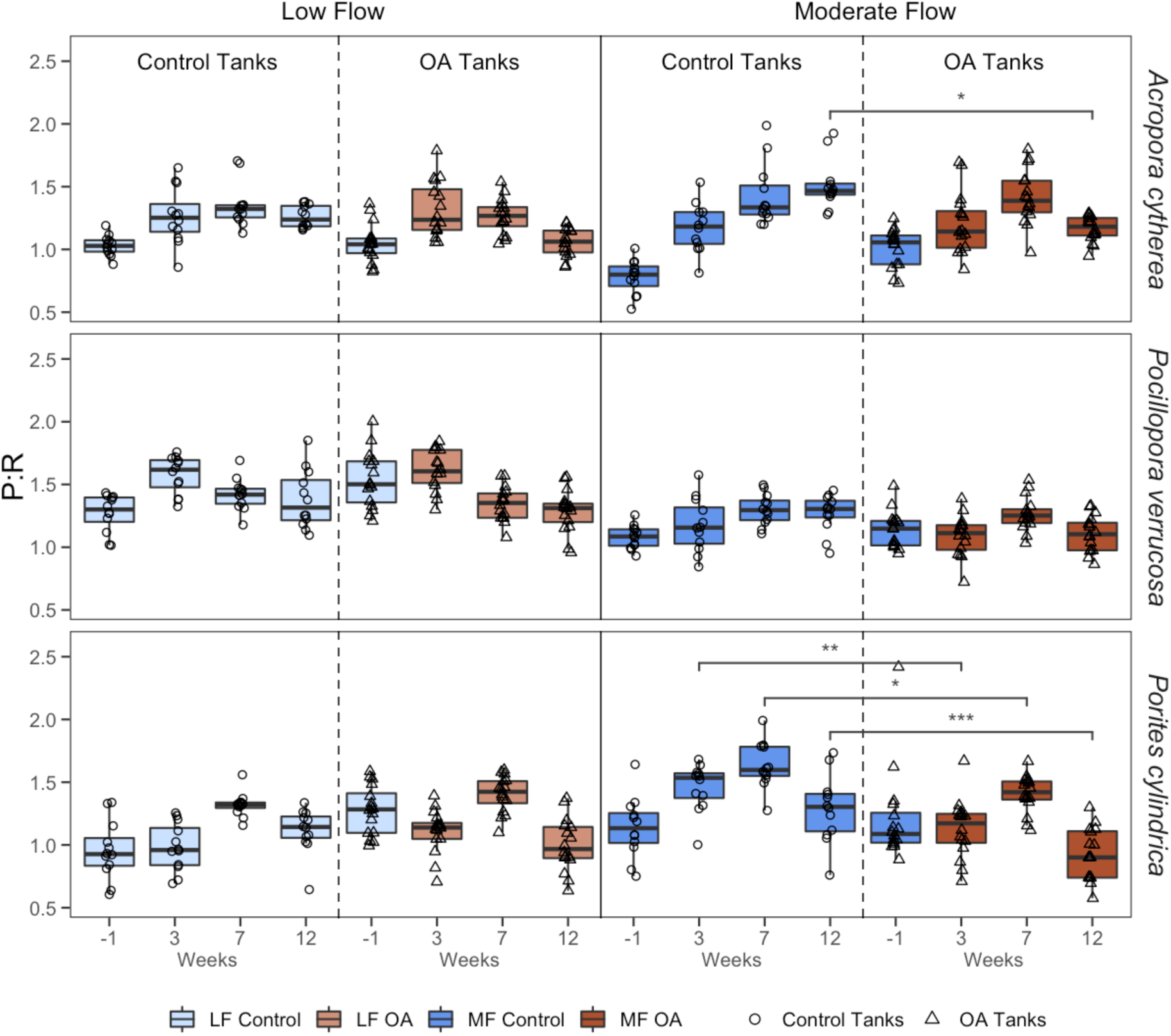
Photosynthesis:Respiration (P:R) ratio of *Acropora cytherea*, *Pocillopora verrucosa*, and *Porites cylindrica* in a control and ocean acidification (OA) treatment and measured under low flow (LF, 2 cm s^-1^) and moderate flow (MF, 6 cm s^-1^) conditions during the acclimation period (measured one week before the start of gradual pH decrease) and experimental period (after three, seven, and 12 weeks under OA, including two weeks of gradual pH decrease). Data from the acclimation period are presented separated by the respective treatment applied during the OA phase. Boxes represent the first and third quartiles with lines as medians and whiskers as the minimum and maximum values or up to the 1.5 × interquartile range (IQR), whichever is reached first. Stars indicate significant differences between the control and OA treatment within each time point and flow condition (p < 0.001***, p < 0.01**, p < 0.05*, from linear mixed-effects models with ANOVA).

## 4. DISCUSSION

Our study shows that the physiological response of corals to future OA conditions varies with water flow conditions. In this study, all coral species showed reduced calcification and surface area growth rates under near-natural OA conditions with diel pH oscillation. In addition, short periods of low water flow modulated P:R ratios, which developed with OA-exposure time and differed among species with potential downstream effects on their calcification rates.

### 4.1. Decreased growth under ocean acidification with diel pH oscillation

In our study, calcification and surface growth decreased in all species under OA, which is consistent with previous findings (e.g., Sekizawa et al. 2017) and the generalised effect of OA on coral calcification (Kornder et al. 2018). Still, calcification in *Acropora cytherea* may be unaffected by OA at stable and even higher *p*CO_2_ than tested here (1,000 µatm *p*CO_2_; Godefroid et al. 2021). Likewise, *Pocillopora verrucosa* maintained stable calcification rates with moderately and highly elevated *p*CO_2_ under stable pH conditions (700 and 1,000 µatm *p*CO_2_; Comeau et al. 2019b). In coral reefs, pH oscillates naturally with diel ranges, which vary between and within reefs (Hannan et al. 2020; Cyronak et al. 2020) and are expected to increase under future OA conditions (Shaw et al. 2013). pH variability is thus an important element of biological OA manipulation experiments that is essential for extrapolating responses observed in the laboratory to coral reefs (Ziegler et al. 2021). In our study, pH was not stable but had a simulated diel oscillation of 0.4 pH units, which is representative of ranges present in some reefs (Cyronak et al. 2020). Since pH oscillation may exacerbate OA-induced calcification decreases (Comeau et al. 2014a), the calcification responses observed in our study are thus potentially associated with diel pH oscillation as observed in natural reef ecosystems.

### 4.2. Physiological response to ocean acidification dependent on exposure time and water flow

OA is a slowly developing press disturbance. Therefore, short-term acute assays will fail to capture the breadth of organismal responses to naturally-occurring acidification. Here, we assessed the progression of OA effects by monitoring P:R ratios over time. P:R ratios were in line with daily P:R ratios reported previously (∼1.1; Jacquemont et al. 2022) and also when calculated as a direct P:R ratio (an estimate of autotrophic capability during daylight hours; ∼2.3; Biscéré et al. 2019). Under OA, P:R ratios decreased at moderate flow, with time-delayed differential responses, except in *P. verrucosa*, where they remained stable. These results could suggest an increasing reliance on heterotrophically fixed energy under OA, consistent with the reported alleviation of OA effects on calcification with increased feeding (Towle et al. 2015). These data thus indicate different vulnerabilities among species with variable trophic dependence.

Specifically, the response of *A. cytherea*, which showed significant differences between treatments only after 12 weeks of exposure to OA, might indicate an initial compensation for adverse effects that the coral was then unable to sustain over time. Acroporids are highly autotrophic, which is a trophic mode associated with high susceptibility to environmental stress, such as high temperature (Conti-Jerpe et al. 2020). However, some *Acropora* spp. have the capacity to increase feeding rates under OA conditions and heterotrophically compensate for OA effects (Towle et al. 2015), which might explain the delayed response observed in *A. cytherea*. Therefore, our results support the classification of *A. cytherea* with other acroporids as OA-susceptible in the long term (Kornder et al. 2018), despite being able to potentially compensate for short-term OA challenges.

With changes in P:R ratios after just three weeks of exposure to OA, the response of *Porites cylindrica* to OA was the most immediate in our study. While massive *Porites* spp. are consistently observed to be OA-resilient (Fabricius et al. 2011; Comeau et al. 2019b), poritids with branching growth forms, such as *P. cylindrica,* appear more vulnerable (Comeau et al. 2014b). However, its overall milder response compared to *A. cytherea* may be due to the mixotrophic strategy of poritids, i.e., their more balanced contributions from autotrophy and heterotrophy (Conti-Jerpe et al. 2020). In contrast to *A. cytherea* and *P. cylindrica*, *P. verrucosa* maintained stable P:R ratios across treatments throughout the experiment and was thus more resilient to OA, supporting the current view that pocilloporids are highly resilient (Kornder et al. 2018). A higher contribution of heterotrophic feeding and physiological plasticity in *Pocillopora* than in *Acropora* or *Porites* spp. may underlie these differences (Hoogenboom et al. 2015; Radice et al. 2019).

Furthermore, our results from *A. cytherea* and *P. cylindrica* confirm the expectation of stronger effects under moderate flow than under low flow. Ecologically this means that some species threatened by OA on a global scale might benefit from local environmental variation and offset OA effects on local scales (e.g., in environments with large short-term OA fluctuations and/or low flow such as reef flats under calm weather; Lowe et al. 2009; Shaw et al. 2012). Accordingly, low-flow environments are considered refuge environments from OA for many calcifying organisms (Hurd 2015). Notably, the effects of heat stress, which is a pulse disturbance, may be stronger for *Acropora* spp. under low flow than high flow conditions (Nakamura and Van Woesik 2001; Page et al. 2021), and recovery from bleaching events may also be faster under high flow conditions (Fifer et al. 2021). Yet, field studies have found lower bleaching intensity at lagoon sites than high-flow environments (McClanahan et al. 2007; Hoogenboom et al. 2017), which may not be the case for all reef habitats (e.g., Ainsworth et al. 2021). Therefore, future coral reef conservation strategies to address climate change will likely benefit from a diverse portfolio of refugia to balance trade-offs associated with reef-specific characteristics and timescales of stressors.

The low-flow effects reported in this study might be underestimated when compared to laminar low-flow conditions. Given our respirometry setup, the flow regimes tested were likely relatively turbulent with lower intensity in the low flow condition due to its lower velocity. Under turbulent flow conditions, boundary layer behaviour may be limited (Reidenbach et al. 2006), reducing the effect of flow changes on coral physiology (Lesser et al. 1994). Therefore, our results support the notion of milder OA effects with reduced flow, even when flow is relatively turbulent, as may occur on coral reefs. The flow conditions simulated in this study could be representative of conditions in reef areas where oscillatory flow may limit the development of boundary layers, such as reef flats (Davis et al. 2021). Still, *in situ* flow regimes are influenced by various factors (Monismith 2007; Davis et al. 2021) and are thus generally more complex than the flow conditions in our study. Also, coral colonies larger than the coral fragments in our study may have a larger effect on the flow patterns around them (e.g., Hench and Rosman 2013; Hossain and Staples 2020). Therefore, future studies incorporating ecologically relevant flow dynamics and a range of colony sizes and shapes will be important to disentangle OA and flow effects.

## 5. CONCLUSIONS

Our study indicates that short periods of low flow modulate the physiological response of corals to OA. We show that OA conditions of moderately elevated *p*CO_2_ and naturally oscillating pH I) reduce coral calcification and surface growth rates and that II) coral species display differential time-delayed P:R ratio responses to OA, which may be mitigated by temporarily reduced flow conditions. This differential progression of OA responses over time may be related to differences in trophic strategy and explain the variable susceptibility to long-term OA among coral species, which should be subject of follow-up work. Future research on this topic could potentially inform the design and management of coral nurseries. Overall, our results highlight that the combination of long-term OA exposure in the variable hydrodynamic conditions of coral reefs may lead to complex biological outcomes that require consideration of the spatial and temporal scales at which they occur. Finally, *in situ* flow regimes are generally more complex than the low flow conditions in our study. Therefore, future studies incorporating ecologically relevant flow regimes and dynamics will be important to disentangle OA and flow effects and better understand the potential of low-flow environments as refugia.

## DATA AVAILABILITY

The datasets and code for the analyses presented in this study can be found in the online Figshare repository and are accessible at https://doi.org/10.6084/m9.figshare.23538474.

## ACKNOWLEDGEMENTS

We thank Giulia Puntin (JLU, Germany) for help during respirometry measurements. This study was conducted as part of the ‘Ocean2100’ global change simulation project of the Colombian-German Center of Excellence in Marine Sciences (CEMarin), funded by the German Academic Exchange Service (DAAD, project number 57480468).

## REFERENCES

Ainsworth TD, Leggat W, Silliman BR, Lantz CA, Bergman JL, Fordyce AJ, Page CE, Renzi JJ, Morton J, Eakin CM, Heron SF (2021) Rebuilding relationships on coral reefs: Coral bleaching knowledge-sharing to aid adaptation planning for reef users. BioEssays. doi: 10.1002/bies.202100048

Anthony KRN, Diaz-Pulido G, Verlinden N, Tilbrook B, Andersson AJ (2013) Benthic buffers and boosters of ocean acidification on coral reefs. Biogeosciences 10:4897–4909. doi: 10.5194/bg-10-4897-2013

Atkinson MJ, Bilger RW (1992) Effects of water velocity on phosphate uptake in coral reef-flat communities. Limnol Oceanogr 37:273–279. doi: 10.4319/lo.1992.37.2.0273

Bates D, Mächler M, Bolker B, Walker S (2015) Fitting linear mixed-effects models using lme4. J Stat Softw 67:1–48. doi: 10.18637/jss.v067.i01

Bedwell-Ivers HE, Koch MS, Peach KE, Joles L, Dutra E, Manfrino C (2017) The role of *in hospite* zooxanthellae photophysiology and reef chemistry on elevated *p*CO_2_ effects in two branching Caribbean corals: *Acropora cervicornis* and *Porites divaricata*. ICES J Mar Sci 74:1103–1112. doi: 10.1093/icesjms/fsw026

Biscéré T, Zampighi M, Lorrain A, Jurriaans S, Foggo A, Houlbrèque F, Rodolfo-Metalpa R (2019) High *p*CO_2_ promotes coral primary production. Biol Lett 15:20180777. doi: 10.1098/rsbl.2018.0777

Cignoni P, Callieri M, Corsini M, Dellepiane M, Ganovelli F, Ranzuglia G (2008) MeshLab: An Open-Source Mesh Processing Tool. Sixth Eurographics Italian Chapter Conference 129–136. doi: 10.2312/LocalChapterEvents/ItalChap/ItalianChapConf2008/129-136

Comeau S, Edmunds PJ, Spindel NB, Carpenter RC (2014a) Diel *p*CO_2_ oscillations modulate the response of the coral *Acropora hyacinthus* to ocean acidification. Mar Ecol Prog Ser 501:99–111. doi: 10.3354/meps10690

Comeau S, Edmunds PJ, Spindel NB, Carpenter RC (2014b) Fast coral reef calcifiers are more sensitive to ocean acidification in short-term laboratory incubations. Limnol Oceanogr 59:1081–1091. doi: 10.4319/lo.2014.59.3.1081

Comeau S, Edmunds PJ, Lantz CA, Carpenter RC (2014c) Water flow modulates the response of coral reef communities to ocean acidification. Sci Rep 4:1–6. doi: 10.1038/srep06681

Comeau S, Cornwall CE, DeCarlo TM, Krieger E, McCulloch MT (2018) Similar controls on calcification under ocean acidification across unrelated coral reef taxa. Glob Chang Biol 24:4857–4868. doi: 10.1111/gcb.14379

Comeau S, Cornwall CE, Pupier CA, DeCarlo TM, Alessi C, Trehern R, McCulloch MT (2019a) Flow-driven micro-scale pH variability affects the physiology of corals and coralline algae under ocean acidification. Sci Rep 9:12829. doi: 10.1038/s41598-019-49044-w

Comeau S, Cornwall CE, DeCarlo TM, Doo SS, Carpenter RC, McCulloch MT (2019b) Resistance to ocean acidification in coral reef taxa is not gained by acclimatization. Nat Clim Chang 9:477–483. doi: 10.1038/s41558-019-0486-9

Conti-Jerpe IE, Thompson PD, Wong CWM, Oliveira NL, Duprey NN, Moynihan MA, Baker DM (2020) Trophic strategy and bleaching resistance in reef-building corals. Sci Adv. doi: 10.1126/sciadv.aaz5443

Cyronak T, Takeshita Y, Courtney TA, DeCarlo EH, Eyre BD, Kline DI, Martz T, Page H, Price NN, Smith J, Stoltenberg L, Tresguerres M, Andersson AJ (2020) Diel temperature and pH variability scale with depth across diverse coral reef habitats. Limnol Oceanogr Lett 5:193–203. doi: 10.1002/lol2.10129

Davis KA, Pawlak G, Monismith SG (2021) Turbulence and coral reefs. Ann Rev Mar Sci 13:343–373. doi: 10.1146/annurev-marine-042120-071823

Dennison WC, Barnes DJ (1988) Effect of water motion on coral photosynthesis and calcification. J Exp Mar Biol Ecol 115:67–77. doi: 10.1016/0022-0981(88)90190-6

Dickson AG, Millero FJ (1987) A comparison of the equilibrium constants for the dissociation of carbonic acid in seawater media. Deep Sea Research Part A Oceanographic Research Papers 34:1733–1743.

Dickson AG, Afghan JD, Anderson GC (2003) Reference materials for oceanic CO_2_ analysis: A method for the certification of total alkalinity. Mar Chem 80:185–197. doi: 10.1016/S0304-4203(02)00133-0

Dickson AG, Sabine CL, Christian JR (2007) Guide to best practices for ocean CO_2_ measurements. PICES Special Publication 3, CA

Doney SC, Fabry VJ, Feely RA, Kleypas JA (2009) Ocean acidification: The other CO_2_ problem. Ann Rev Mar Sci 1:169–192. doi: 10.1146/annurev.marine.010908.163834

Enochs IC, Manzello DP, Jones PJ, Aguilar C, Cohen K, Valentino L, Schopmeyer S, Kolodziej G, Jankulak M, Lirman D (2018) The influence of diel carbonate chemistry fluctuations on the calcification rate of *Acropora cervicornis* under present day and future acidification conditions. J Exp Mar Biol Ecol 506:135–143. doi: 10.1016/j.jembe.2018.06.007

Fabricius KE, Langdon C, Uthicke S, Humphrey C, Noonan S, De’ath G, Okazaki R, Muehllehner N, Glas MS, Lough JM (2011) Losers and winners in coral reefs acclimatized to elevated carbon dioxide concentrations. Nat Clim Chang 1:165–169. doi: 10.1038/nclimate1122

Fifer J, Bentlage B, Lemer S, Fujimura AG, Sweet M, Raymundo LJ (2021) Going with the flow: How corals in high-flow environments can beat the heat. Mol Ecol 30:2009–2024. doi: 10.1111/mec.15869

Fox J, Weisberg S (2019) An R companion to applied regression, Third. Sage, Thousand Oaks, CA

Godefroid M, Arçuby R, Lacube Y, Espiau B, Dupont S, Gazeau F, Metian M, Hédouin L (2021) More than local adaptation: High diversity of response to seawater acidification in seven coral species from the same assemblage in French Polynesia. J Mar Biol Assoc United Kingdom 101:675–683. doi: 10.1017/S0025315421000618

Godefroid M, Dupont S, Metian M, Hédouin L (2022) Two decades of seawater acidification experiments on tropical scleractinian corals: Overview, meta-analysis and perspectives. Mar Pollut Bull 178:113552. doi: 10.1016/j.marpolbul.2022.113552

Green RH, Lowe RJ, Buckley ML (2018) Hydrodynamics of a tidally forced coral reef atoll. J Geophys Res Oceans 123:7084–7101. doi: 10.1029/2018JC013946

Hannan KD, Miller GM, Watson S-A, Rummer JL, Fabricius K, Munday PL (2020) Diel *p*CO_2_ variation among coral reefs and microhabitats at Lizard Island, Great Barrier Reef. Coral Reefs 39:1391–1406. doi: 10.1007/s00338-020-01973-z

Hector A, von Felten S, Schmid B (2010) Analysis of variance with unbalanced data: An update for ecology & evolution. J Anim Ecol 79:308–316. doi: 10.1111/j.1365-2656.2009.01634.x

Hench JL, Rosman JH (2013) Observations of spatial flow patterns at the coral colony scale on a shallow reef flat. J Geophys Res Oceans 118:1142–1156. doi: 10.1002/jgrc.20105

Hench JL, Leichter JJ, Monismith SG (2008) Episodic circulation and exchange in a wave-driven coral reef and lagoon system. Limnol Oceanogr 53:2681–2694. doi: 10.4319/lo.2008.53.6.2681

Hoogenboom M, Rottier C, Sikorski S, Ferrier-Pagès C (2015) Among-species variation in the energy budgets of reef-building corals: Scaling from coral polyps to communities. J Exp Biol 218:3866–3877. doi: 10.1242/jeb.124396

Hoogenboom MO, Frank GE, Chase TJ, Jurriaans S, Álvarez-Noriega M, Peterson K, Critchell K, Berry KLE, Nicolet KJ, Ramsby B, Paley AS (2017) Environmental drivers of variation in bleaching severity of *Acropora* species during an extreme thermal anomaly. Front Mar Sci. doi: 10.3389/fmars.2017.00376

Hossain MM, Staples AE (2020) Effects of coral colony morphology on turbulent flow dynamics. PLoS One 15:1–25. doi: 10.1371/journal.pone.0225676

Hurd CL (2015) Slow-flow habitats as refugia for coastal calcifiers from ocean acidification. J Phycol 51:599–605. doi: 10.1111/jpy.12307

IPCC (2023) Atlas. In: o (eds) Climate Change 2021 – The Physical Science Basis: Working Group I Contribution to the Sixth Assessment Report of the Intergovernmental Panel on Climate Change. Cambridge University Press, Cambridge, UK and New York, NY, USA, pp 1927–2058

Iturbide M, Fernández J, Gutiérrez JM, Pirani A, Huard D, Al Khourdajie A, Baño-Medina J, Bedia J, Casanueva A, Cimadevilla E, Cofiño AS, De Felice M, Diez-Sierra J, García-Díez M, Goldie J, Herrera DA, Herrera S, Manzanas R, Milovac J, Radhakrishnan A, San-Martín D, Spinuso A, Thyng KM, Trenham C, Yelekçi Ö (2022) Implementation of FAIR principles in the IPCC: The WGI AR6 Atlas repository. Sci Data 9:629. doi: 10.1038/s41597-022-01739-y

Jacquemont J, Houlbrèque F, Tanvet C, Rodolfo-Metalpa R (2022) Long-term exposure to an extreme environment induces species-specific responses in corals’ photosynthesis and respiration rates. Mar Biol 169:82. doi: 10.1007/s00227-022-04063-6

Jokiel PL, Maragos JE, Franzisket L (1978) Coral growth: Buoyant weight technique. In: Stoddart DR, Johannes RE (eds) Coral reefs: research methods. UNESCO, pp 529– 542

Kornder NA, Riegl BM, Figueiredo J (2018) Thresholds and drivers of coral calcification responses to climate change. Glob Chang Biol 24:5084–5095. doi: 10.1111/gcb.14431

Kroeker KJ, Kordas RL, Crim R, Hendriks IE, Ramajo L, Singh GS, Duarte CM, Gattuso J (2013) Impacts of ocean acidification on marine organisms: Quantifying sensitivities and interaction with warming. Glob Chang Biol 19:1884–1896. doi: 10.1111/gcb.12179

Langsrud Ø (2003) ANOVA for unbalanced data: Use Type II instead of Type III sums of squares. Stat Comput 13:163–167. doi: 10.1023/A:1023260610025

Lenth R V. (2021) emmeans: Estimated Marginal Means, aka Least-Squares Means. R package version 1.6.2–1. https://cran.r-project.org/package=emmeans.

Lesser MP, Weis VM, Patterson MR, Jokiel PL (1994) Effects of morphology and water motion on carbon delivery and productivity in the reef coral, *Pocillopora damicornis* (Linnaeus): Diffusion barriers, inorganic carbon limitation, and biochemical plasticity. J Exp Mar Biol Ecol 178:153–179. doi: 10.1016/0022-0981(94)90034-5

Lindhart M, Rogers JS, Maticka SA, Woodson CB, Monismith SG (2021) Wave modulation of flows on open and closed reefs. J Geophys Res Oceans 126:e2020JC016645. doi: 10.1029/2020JC016645

Lowe RJ, Falter JL (2015) Oceanic forcing of coral reefs. Ann Rev Mar Sci 7:43–66. doi: 10.1146/annurev-marine-010814-015834

Lowe RJ, Falter JL, Monismith SG, Atkinson MJ (2009) A numerical study of circulation in a coastal reef-lagoon system. J Geophys Res 114:C06022. doi: 10.1029/2008JC005081

Lüdecke D (2021) sjPlot: Data visualization for statistics in social science. R package version 2.8.9. https://CRAN.R-project.org/package=sjPlot.

Lüdecke D, Ben-Shachar M, Patil I, Waggoner P, Makowski D (2021) performance: An R package for assessment, comparison and testing of statistical models. J Open Source Softw 6:3139. doi: 10.21105/joss.03139

Madin JS, Black KP, Connolly SR (2006) Scaling water motion on coral reefs: From regional to organismal scales. Coral Reefs 25:635–644. doi: 10.1007/s00338-006-0137-2

McClanahan TR, Ateweberhan M, Muhando CA, Maina J, Mohammed MS (2007) Effects of climate and seawater temperature variation on coral bleaching and mortality. Ecol Monogr 77:503–525. doi: 10.1890/06-1182.1

McCloskey LR, Wethey DS, Porter JW (1978) Measurement and interpretation of photosynthesis and respiration in reef corals. In: Stoddart DR, Johannes RE (eds) Coral reefs: research methods. UNESCO, pp 379–396

McLachlan RH, Price JT, Muñoz-Garcia A, Weisleder NL, Jury CP, Toonen RJ, Grottoli AG (2021) Environmental gradients drive physiological variation in Hawaiian corals. Coral Reefs 40:1505–1523. doi: 10.1007/s00338-021-02140-8

Mehrbach C, Culberson CH, Hawley JE, Pytkowicx RM (1973) Measurement of the apparent dissociation constants of carbonic acid in seawater at atmospheric pressure. Limnol Oceanogr 18:897–907.

Millero FJ (2013) Chemical oceanography, 4th edn. CRC Press, Boca Raton, Florida

Monismith SG (2007) Hydrodynamics of coral reefs. Annu Rev Fluid Mech 39:37–55. doi: 10.1146/annurev.fluid.38.050304.092125

Nakamura T, Van Woesik R (2001) Water-flow rates and passive diffusion partially explain differential survival of corals during the 1998 bleaching event. Mar Ecol Prog Ser 212:301–304. doi: 10.3354/meps212301

Noisette F, Pansch C, Wall M, Wahl M, Hurd CL (2022) Role of hydrodynamics in shaping chemical habitats and modulating the responses of coastal benthic systems to ocean global change. Glob Chang Biol 28:3812–3829. doi: 10.1111/gcb.16165

Osinga R, Derksen-Hooijberg M, Wijgerde T, Verreth JAJ (2017) Interactive effects of oxygen, carbon dioxide and flow on photosynthesis and respiration in the scleractinian coral *Galaxea fascicularis*. J Exp Biol 220:2236–2242. doi: 10.1242/jeb.140509

Page CE, Leggat W, Heron SF, Fordyce AJ, Ainsworth TD (2021) High flow conditions mediate damaging impacts of sub-lethal thermal stress on corals’ endosymbiotic algae. Conserv Physiol 9:1–19. doi: 10.1093/conphys/coab046

Patterson MR, Sebens KP, Olson RR (1991) In situ measurements of flow effects on primary production and dark respiration in reef corals. Limnol Oceanogr 36:936–948. doi: 10.4319/lo.1991.36.5.0936

Pelletier G, Lewis E, Wallace D (2007) CO_2_Sys. xls: A calculator for the CO_2_ system in seawater for Microsoft Excel/VBA. Washington State Department of Ecology/Brookhaven National Laboratory, Olympia, WA/Upton, NY, USA

R Core Team (2021) R: A language and environment for statistical computing. R Foundation for Statistical Computing, Vienna, Austria

Rades M, Schubert P, Ziegler M, Kröckel M, Reichert J (2022) Building plan for a temperature-controlled multi-point stirring incubator. protocols.io. doi: 10.17504/protocols.io.dm6gpb34dlzp/v1

Radice VZ, Brett MT, Fry B, Fox MD, Hoegh-Guldberg O, Dove SG (2019) Evaluating coral trophic strategies using fatty acid composition and indices. PLoS One 14:e0222327. doi: 10.1371/journal.pone.0222327

Reichert J, Schellenberg J, Schubert P, Wilke T (2016) 3D scanning as a highly precise, reproducible, and minimally invasive method for surface area and volume measurements of scleractinian corals. Limnol Oceanogr Methods 14:518–526. doi: 10.1002/lom3.10109

Reidenbach MA, Koseff JR, Monismith SG, Steinbuckc J V., Genin A (2006) The effects of waves and morphology on mass transfer within branched reef corals. Limnol Oceanogr 51:1134–1141. doi: 10.4319/lo.2006.51.2.1134

Rex A, Montebon F, Yap HT (1995) Metabolic responses of the scleractinian coral *Porites cylindrica* Dana to water motion. I. Oxygen flux studies. J Exp Mar Biol Ecol 186:33–52. doi: 10.1016/0022-0981(95)00150-P

Reynaud S, Leclercq N, Romaine-Lioud S, Ferrier-Pagés C, Jaubert J, Gattuso J (2003) Interacting effects of CO_2_ partial pressure and temperature on photosynthesis and calcification in a scleractinian coral. Glob Chang Biol 9:1660–1668. doi: 10.1046/j.1529-8817.2003.00678.x

Roik A, Röthig T, Roder C, Ziegler M, Kremb SG, Voolstra CR (2016) Year-long monitoring of physico-chemical and biological variables provide a comparative baseline of coral reef functioning in the central Red Sea. PLoS One 11:e0163939. doi: 10.1371/journal.pone.0163939

RStudio Team (2021) RStudio: Integrated development environment for R.

RStudio, PBC, Boston, MA Schutter M, Crocker J, Paijmans A, Janse M, Osinga R, Verreth AJ, Wijffels RH (2010) The effect of different flow regimes on the growth and metabolic rates of the scleractinian coral *Galaxea fascicularis*. Coral Reefs 29:737–748. doi: 10.1007/s00338-010-0617-2

Sekizawa A, Uechi H, Iguchi A, Nakamura T, Kumagai NH, Suzuki A, Sakai K, Nojiri Y (2017) Intraspecific variations in responses to ocean acidification in two branching coral species. Mar Pollut Bull 122:282–287. doi: 10.1016/j.marpolbul.2017.06.061

Shashar N, Cohen Y, Loya Y (1993) Extreme diel fluctuations of oxygen in diffusive boundary layers surrounding stony corals. Biol Bull 185:455–461. doi: 10.2307/1542485

Shaw EC, McNeil BI, Tilbrook B (2012) Impacts of ocean acidification in naturally variable coral reef flat ecosystems. J Geophys Res Oceans 117:C03038. doi: 10.1029/2011JC007655

Shaw EC, McNeil BI, Tilbrook B, Matear R, Bates ML (2013) Anthropogenic changes to seawater buffer capacity combined with natural reef metabolism induce extreme future coral reef CO_2_ conditions. Glob Chang Biol 19:1632–1641. doi: 10.1111/gcb.12154

Sheppard CRC, Davy SK, Pilling GM, Graham NAJ (2018) The abiotic environment. In: Sheppard C, Davy S, Pilling G, Graham N (eds) The Biology of Coral Reefs, 2nd edn. Oxford University Press, Oxford, UK, pp 68–99

Smith LW, Birkeland C (2007) Effects of intermittent flow and irradiance level on back reef *Porites* corals at elevated seawater temperatures. J Exp Mar Biol Ecol 341:282–294. doi: 10.1016/j.jembe.2006.10.053

Spencer Davies P (1989) Short-term growth measurements of corals using an accurate buoyant weighing technique. Mar Biol 101:389–395. doi: 10.1007/BF00428135

Takahashi T (1982) Carbonate chemistry. In: GEOSECS Pacific expedition, Volume 3, Hydrographic Data 1973-1974. National Science Foundation, Washington, DC, pp 77– 83

Towle EK, Enochs IC, Langdon C (2015) Threatened Caribbean coral is able to mitigate the adverse effects of ocean acidification on calcification by increasing feeding rate. PLoS One 10:e0123394. doi: 10.1371/journal.pone.0123394

UNEP-WCMC, WorldFish Centre, WRI, TNC (2021) Global distribution of warm-water coral reefs, compiled from multiple sources including the Millennium Coral Reef Mapping Project. Version 4.1. Includes contributions from IMaRS-USF and IRD (2005), IMaRS-USF (2005) and Spalding et al. (2001). In: UN Environment World Conservation Monitoring Centre. Cambridge, UK. 10.34892/t2wk-5t34. Accessed 22 Jul 2023

Watson S-A, Fabricius KE, Munday PL (2017) Quantifying *p*CO_2_ in biological ocean acidification experiments: A comparison of four methods. PLoS One 12:e0185469. doi: 10.1371/journal.pone.0185469

Wickham H (2016) ggplot2: Elegant graphics for data analysis. Springer-Verlag, New York

Ziegler M, Anton A, Klein SG, Rädecker N, Geraldi NR, Schmidt-Roach S, Saderne V, Mumby PJ, Cziesielski MJ, Martin C, Frölicher TL, Pandolfi JM, Suggett DJ, Aranda M, Duarte CM, Voolstra CR (2021) Integrating environmental variability to broaden the research on coral responses to future ocean conditions. Glob Chang Biol 27:5532–5546. doi: 10.1111/gcb.15840

