## Supplementary Material for "Short periods of decreased water flow may modulate long-term ocean acidification in reef-building corals"

### Supplementary Text

#### Calculation of photosynthesis and respiration rates

Photosynthesis and respiration were calculated as the change in dissolved oxygen per incubation volume (calculated as the difference between water volume and volume of the coral fragment with its tile) and normalised to incubation time and surface area of the coral fragment. We used the surface area measured at $t_{-4}$ and $t_{14}$ to surface-normalise respiration and photosynthesis measured at $t_{-1}$ and $t_{12}$, respectively. Respiration and photosynthesis rates were not calculated for $t_3$ and $t_7$ because no surface measurements were performed during these time points for surface normalisation. Gross photosynthesis was calculated as the sum of net photosynthesis and respiration values.

### Supplementary Tables

**Table S1** Details of coral species used in the experiment.

| Species | Number of colonies | Origin | Collection year | CITES number |
|---|---|---|---|---|
| *Acropora cytherea* | 4 | Saudi Arabia (Red Sea) | 2019 | 19-SA-000089-PD |
| *Pocillopora verrucosa* | 2 | Saudi Arabia (Red Sea) | 2015 | 15-SA-000882-PD |
| | 2 | Indonesia (Indo-Pacific) | 2007 | 14846/IV/SATS-LN/2007 |
| *Porites cylindrica* | 4 | Indonesia (Indo-Pacific) | 2017 | 17nl241924/11 |

**Table S2** Brands of artificial-seawater salt used alternately during the acclimation and experimental period in the 'Ocean2100' experimental facility.

| Salt | Brand |
|---|---|
| Deep Blue Sea-Salt | AquaPerfekt, Germany |
| Instant Ocean | Aquarium Systems, France |
| Reef Crystals | Aquarium Systems, France |
| Sea Salt | Aquaforest, Poland |

**Table S3** Respiration, net photosynthesis, and gross photosynthesis of *Acropora cytherea*, *Pocillopora verrucosa*, and *Porites cylindrica* in a control and ocean acidification (OA) treatment and measured in low and moderate flow (2 and 6 cm $s^{-1}$, respectively). Values are means ± SD (in µg $O_2$ $cm^{-2}$ $h^{-1}$) of initial ($t_{-1}$, one week before the start of gradual pH decrease) and final rates ($t_{12}$, after 12 weeks under OA conditions, including two weeks of gradual pH decrease). Sample size (*n*) is provided and 'pre-OA' refers to the experimental OA group before the start of the OA treatment, with conditions equivalent to the control treatment.

| Species | Time Point | Flow | Treatment | Respiration | Net Photosynthesis | Gross Photosynthesis | *n* |
|---|---|---|---|---|---|---|---|
| *Acropora cytherea* | $t_{-1}$ | Low | Control | 19.2 ± 3.8 | 23.8 ± 5.1 | 43.0 ± 8.3 | 12 |
| *Acropora cytherea* | $t_{-1}$ | Low | pre-OA | 17.5 ± 3.4 | 22.3 ± 6.4 | 39.8 ± 8.5 | 16 |
| *Acropora cytherea* | $t_{-1}$ | Moderate | Control | 23.3 ± 5.2 | 16.0 ± 6.9 | 39.3 ± 9.9 | 12 |
| *Acropora cytherea* | $t_{-1}$ | Moderate | pre-OA | 18.4 ± 3.8 | 21.6 ± 5.7 | 40.0 ± 7.3 | 16 |
| *Acropora cytherea* | $t_{12}$ | Low | Control | 14.9 ± 4.6 | 26.1 ± 8.9 | 41.0 ± 13.3 | 12 |
| *Acropora cytherea* | $t_{12}$ | Low | OA | 12.1 ± 3.1 | 15.6 ± 4.3 | 27.7 ± 6.8 | 16 |
| *Acropora cytherea* | $t_{12}$ | Moderate | Control | 9.8 ± 2.2 | 22.4 ± 6.7 | 32.2 ± 8.5 | 12 |
| *Acropora cytherea* | $t_{12}$ | Moderate | OA | 9.1 ± 2.0 | 13.8 ± 3.1 | 22.9 ± 4.8 | 16 |
| *Pocillopora verrucosa* | $t_{-1}$ | Low | Control | 20.2 ± 3.5 | 35.8 ± 8.0 | 56.0 ± 10.3 | 12 |
| *Pocillopora verrucosa* | $t_{-1}$ | Low | pre-OA | 16.3 ± 3.1 | 37.4 ± 6.6 | 53.7 ± 8.4 | 16 |
| *Pocillopora verrucosa* | $t_{-1}$ | Moderate | Control | 26.1 ± 5.1 | 35.4 ± 8.6 | 61.5 ± 12.8 | 12 |
| *Pocillopora verrucosa* | $t_{-1}$ | Moderate | pre-OA | 21.0 ± 5.2 | 30.6 ± 7.3 | 51.6 ± 11.4 | 16 |
| *Pocillopora verrucosa* | $t_{12}$ | Low | Control | 9.6 ± 1.3 | 19.0 ± 4.0 | 28.7 ± 4.2 | 12 |
| *Pocillopora verrucosa* | $t_{12}$ | Low | OA | 9.4 ± 1.1 | 17.0 ± 3.9 | 26.4 ± 4.4 | 16 |
| *Pocillopora verrucosa* | $t_{12}$ | Moderate | Control | 10.5 ± 1.1 | 18.5 ± 3.9 | 29.0 ± 4.6 | 12 |
| *Pocillopora verrucosa* | $t_{12}$ | Moderate | OA | 10.1 ± 1.1 | 14.4 ± 3.8 | 24.5 ± 4.4 | 16 |
| *Porites cylindrica* | $t_{-1}$ | Low | Control | 29.8 ± 7.6 | 32.0 ± 14.7 | 61.7 ± 18.7 | 12 |
| *Porites cylindrica* | $t_{-1}$ | Low | pre-OA | 15.9 ± 4.5 | 27.9 ± 7.6 | 43.8 ± 10.9 | 16 |
| *Porites cylindrica* | $t_{-1}$ | Moderate | Control | 25.8 ± 7.0 | 36.3 ± 10.5 | 62.1 ± 13.7 | 12 |
| *Porites cylindrica* | $t_{-1}$ | Moderate | pre-OA | 17.5 ± 6.1 | 25.8 ± 7.8 | 43.3 ± 12.7 | 16 |
| *Porites cylindrica* | $t_{12}$ | Low | Control | 8.4 ± 1.5 | 12.2 ± 4.1 | 20.6 ± 5.2 | 12 |
| *Porites cylindrica* | $t_{12}$ | Low | OA | 8.6 ± 1.2 | 10.3 ± 4.2 | 18.9 ± 4.9 | 16 |
| *Porites cylindrica* | $t_{12}$ | Moderate | Control | 9.0 ± 2.0 | 16.0 ± 5.4 | 25.1 ± 6.5 | 12 |
| *Porites cylindrica* | $t_{12}$ | Moderate | OA | 9.2 ± 1.4 | 9.4 ± 4.4 | 18.5 ± 5.1 | 16 |

**Table S4** Numerical output of linear mixed-effects models (LMMs) of calcification (C) and surface growth (Gs). Global models were constructed with species (3 levels: *Acropora cytherea* [Acy], *Pocillopora verrucosa* [Pve], and *Porites cylindrica* [Pcy]) as a fixed factor, and colony and treatment as random factors. Individual species' models were constructed with treatment (2 levels: control and ocean acidification [OA]) as a fixed factor and colony as a random factor. Model formulas are specified. $\sigma^2$, residual variance; $\tau_{00}$, random intercept variance; ICC, intra-class correlation coefficient; N, number of levels of random effects groups; Marginal $R^2$, variance explained by the fixed effects; Conditional $R^2$, variance explained by the entire model

| | | Calcification | | Surface Growth | |
|---|---|---|---|---|---|
| | | C ~ Species + (1 \| Treatment) + (1 \| Colony) | | Gs ~ Species + (1 \| Treatment) + (1 \| Colony) | |
| | Fixed Effects | Estimates | SE | Estimates | SE |
| All Species | (Intercept) | 0.56 | 0.22 | 0.48 | 0.16 |
| | species [Pve] | 1.33 | 0.24 | 0.83 | 0.15 |
| | species [Pcy] | 0.99 | 0.24 | 0.66 | 0.15 |
| | Random Effects | | | | |
| | $\sigma^2$ | 0.09 | | 0.05 | |
| | $\tau_{00}$ Treatment | 0.04 | | 0.03 | |
| | $\tau_{00}$ Colony | 0.10 | | 0.04 | |
| | ICC | 0.60 | | 0.59 | |
| | N Treatment | 2 | | 2 | |
| | N Colony | 12 | | 12 | |
| | Observations | 83 | | 83 | |
| | Marginal $R^2$ / Conditional $R^2$ | 0.582 / 0.833 | | 0.527 / 0.805 | |
| | | C ~ Treatment + (1 \| Colony) | | Gs ~ Treatment + (1 \| Colony) | |
| | Fixed Effects | Estimates | SE | Estimates | SE |
| *Acropora cytherea* | (Intercept) | 0.63 | 0.14 | 0.56 | 0.14 |
| | treatment [OA] | -0.16 | 0.04 | -0.17 | 0.05 |
| | Random Effects | | | | |
| | $\sigma^2$ | 0.01 | | 0.02 | |
| | $\tau_{00}$ Colony | 0.07 | | 0.07 | |
| | ICC | 0.86 | | 0.79 | |
| | N Colony | 4 | | 4 | |
| | Observations | 28 | | 28 | |
| | Marginal $R^2$ / Conditional $R^2$ | 0.069 / 0.868 | | 0.078 / 0.810 | |
| *Pocillopora verrucosa* | (Intercept) | 2.10 | 0.17 | 1.49 | 0.07 |
| | treatment [OA] | -0.41 | 0.16 | -0.34 | 0.09 |
| | Random Effects | | | | |
| | $\sigma^2$ | 0.17 | | 0.06 | |
| | $\tau_{00}$ Colony | 0.06 | | < 0.01 | |
| | ICC | 0.24 | | 0.02 | |
| | N Colony | 4 | | 4 | |
| | Observations | 28 | | 28 | |
| | Marginal $R^2$ / Conditional $R^2$ | 0.159 / 0.365 | | 0.333 / 0.349 | |
| *Porites cylindrica* | (Intercept) | 1.71 | 0.22 | 1.26 | 0.13 |
| | treatment [OA] | -0.32 | 0.12 | -0.25 | 0.10 |
| | Random Effects | | | | |
| | $\sigma^2$ | 0.09 | | 0.07 | |
| | $\tau_{00}$ Colony | 0.17 | | 0.04 | |
| | ICC | 0.65 | | 0.37 | |
| | N Colony | 4 | | 4 | |

| | | | |
|---|---|---|---|
| | Observations | 27 | 27 |
| | Marginal $R^2$ / Conditional $R^2$ | 0.090 / 0.678 | 0.123 / 0.450 |

**Table S5** Numerical output of linear mixed-effects models (LMMs) of photosynthesis:respiration ratio (P:R). Global models were constructed with species (3 levels: *Acropora cytherea* [Acy], *Pocillopora verrucosa* [Pve], and *Porites cylindrica* [Pcy]) as fixed factor; and coral fragment identity (ID), colony, and treatment as random factors. Individual species' models were constructed with treatment (2 levels: control and OA), flow (2 levels: low and moderate), and time (3 levels: $t_3$, $t_7$, $t_{12}$) as fixed factors in a fully crossed design, and ID, colony, and tank as random factors. Model formulas are specified. $\sigma^2$, residual variance; $\tau_{00}$, random intercept variance; ICC, intra-class correlation coefficient; N, number of levels of random effects groups; Marginal $R^2$, variance explained by the fixed effects; Conditional $R^2$, variance explained by the entire model; $t_7$, after seven weeks under OA, including two weeks of gradual pH decrease; $t_{12}$, after 12 weeks

| | | P:R ~ Species + (1 \| Treatment) + (1 \| Coral ID) + (1 \| Colony) + (1 \| Tank) | |
|---|---|---|---|
| | Fixed Effects | Estimates | SE |
| All Species | (Intercept) | 1.28 | 0.05 |
| | species [Pve] | 0.04 | 0.03 |
| | species [Pcy] | -0.06 | 0.03 |
| | Random Effects | | |
| | $\sigma^2$ | | 0.05 |
| | $\tau_{00\ Coral\ ID}$ | | < 0.01 |
| | $\tau_{00\ Colony}$ | | < 0.01 |
| | $\tau_{00\ Tank}$ | | 0.01 |
| | $\tau_{00\ Treatment}$ | | < 0.01 |
| | $N_{Treatment}$ | | 2 |
| | $N_{Coral\ ID}$ | | 84 |
| | $N_{Colony}$ | | 12 |
| | $N_{Tank}$ | | 7 |
| | Observations | | 504 |
| | Marginal $R^2$ / Conditional $R^2$ | | 0.024 / 0.173 |
| | | P:R ~ Treatment * Flow * Time + (1 \| Coral ID) + (1 \| Colony) + (1 \| Tank) | |
| | Fixed Effects | Estimates | SE |
| *Acropora cytherea* | (Intercept) | 1.26 | 0.07 |
| | treatment [OA] | 0.05 | 0.09 |
| | flow [moderate] | -0.09 | 0.07 |
| | timepoint [$t_7$] | 0.09 | 0.07 |
| | timepoint [$t_{12}$] | -0.01 | 0.07 |
| | treatment [OA] * flow [moderate] | -0.02 | 0.09 |
| | treatment [OA] * timepoint [$t_7$] | -0.14 | 0.09 |
| | treatment [OA] * timepoint [$t_{12}$] | -0.25 | 0.09 |
| | flow [moderate] * timepoint [$t_7$] | 0.17 | 0.09 |
| | flow [moderate] * timepoint [$t_{12}$] | 0.35 | 0.09 |
| | (treatment [OA] * flow [moderate]) * timepoint [$t_7$] | 0.10 | 0.12 |
| | (treatment [OA] * flow [moderate]) * timepoint [$t_{12}$] | -0.13 | 0.12 |
| | Random Effects | | |
| | $\sigma^2$ | | 0.03 |
| | $\tau_{00\ Coral\ ID}$ | | < 0.01 |
| | $\tau_{00\ Tank}$ | | 0.01 |
| | $\tau_{00\ Colony}$ | | < 0.01 |
| | ICC | | 0.28 |
| | $N_{Coral\ ID}$ | | 28 |
| | $N_{Colony}$ | | 4 |
| | $N_{Tank}$ | | 7 |
| | Observations | | 168 |
| | Marginal $R^2$ / Conditional $R^2$ | | 0.294 / 0.489 |
| *Pocillopora verrucosa* | (Intercept) | 1.58 | 0.07 |
| | treatment [OA] | 0.04 | 0.08 |
| | flow [moderate] | -0.40 | 0.05 |

| | | | |
|---|---|---|---|
| | timepoint [$t_7$] | -0.15 | 0.05 |
| | timepoint [$t_{12}$] | -0.19 | 0.05 |
| | treatment [OA] * flow [moderate] | -0.12 | 0.07 |
| | treatment [OA] * timepoint [$t_7$] | -0.12 | 0.07 |
| | treatment [OA] * timepoint [$t_{12}$] | -0.14 | 0.07 |
| | flow [moderate] * timepoint [$t_7$] | 0.28 | 0.07 |
| | flow [moderate] * timepoint [$t_{12}$] | 0.29 | 0.07 |
| | (treatment [OA] * flow [moderate]) * timepoint [$t_7$] | 0.17 | 0.10 |
| | (treatment [OA] * flow [moderate]) * timepoint [$t_{12}$] | 0.06 | 0.10 |
| | Random Effects | | |
| | $\sigma^2$ | | 0.02 |
| | $\tau_{00\ Coral\ ID}$ | | < 0.01 |
| | $\tau_{00\ Tank}$ | | 0.01 |
| | $\tau_{00\ Colony}$ | | < 0.01 |
| | ICC | | 0.42 |
| | $N_{Coral\ ID}$ | | 28 |
| | $N_{Colony}$ | | 4 |
| | $N_{Tank}$ | | 7 |
| | Observations | | 168 |
| | Marginal $R^2$ / Conditional $R^2$ | | 0.481 / 0.700 |
| *Porites cylindrica* | (Intercept) | 0.98 | 0.06 |
| | treatment [OA] | 0.13 | 0.08 |
| | flow [moderate] | 0.48 | 0.07 |
| | timepoint [$t_7$] | 0.34 | 0.07 |
| | timepoint [$t_{12}$] | 0.13 | 0.07 |
| | treatment [OA] * flow [moderate] | -0.46 | 0.10 |
| | treatment [OA] * timepoint [$t_7$] | -0.04 | 0.10 |
| | treatment [OA] * timepoint [$t_{12}$] | -0.24 | 0.10 |
| | flow [moderate] * timepoint [$t_7$] | -0.16 | 0.10 |
| | flow [moderate] * timepoint [$t_{12}$] | -0.30 | 0.10 |
| | (treatment [OA] * flow [moderate]) * timepoint [$t_7$] | 0.15 | 0.14 |
| | (treatment [OA] * flow [moderate]) * timepoint [$t_{12}$] | 0.21 | 0.14 |
| | Random Effects | | |
| | $\sigma^2$ | | 0.03 |
| | $\tau_{00\ Coral\ ID}$ | | < 0.01 |
| | $\tau_{00\ Tank}$ | | < 0.01 |
| | $\tau_{00\ Colony}$ | | < 0.01 |
| | ICC | | 0.11 |
| | $N_{Coral\ ID}$ | | 28 |
| | $N_{Colony}$ | | 4 |
| | $N_{Tank}$ | | 7 |
| | Observations | | 168 |
| | Marginal $R^2$ / Conditional $R^2$ | | 0.544 / 0.594 |

**Table S6** Summary of the complete recording of temperature and pH during the experiment. Values are expressed as mean ± SD with measurement replication (*n*). OA, ocean acidification; $pH_T$, pH on the total scale

| Tank | Treatment | Temperature (ºC) | $pH_T$ | Daily Minimum $pH_T$ | Daily Maximum $pH_T$ | Daily Range $pH_T$ |
|---|---|---|---|---|---|---|
| 1 | Control | 25.9 ± 0.2 (3,988) | 7.95 ± 0.12 (3,873) | 7.77 ± 0.06 (84) | 8.13 ± 0.06 (84) | 0.36 ± 0.05 (84) |
| 2 | OA | 25.9 ± 0.2 (3,606) | 7.76 ± 0.13 (3,709) | 7.58 ± 0.07 (84) | 7.94 ± 0.07 (84) | 0.36 ± 0.07 (84) |
| 3 | Control | 25.9 ± 0.2 (3,963) | 7.97 ± 0.12 (3,859) | 7.78 ± 0.05 (84) | 8.15 ± 0.05 (84) | 0.36 ± 0.06 (84) |
| 4 | OA | 25.9 ± 0.2 (3,771) | 7.77 ± 0.13 (3,830) | 7.59 ± 0.07 (84) | 7.96 ± 0.06 (84) | 0.37 ± 0.08 (84) |
| 5 | Control | 26.0 ± 0.3 (3,413) | 7.98 ± 0.13 (3,785) | 7.80 ± 0.06 (84) | 8.17 ± 0.07 (84) | 0.36 ± 0.06 (84) |
| 6 | OA | 26.0 ± 0.2 (3,408) | 7.78 ± 0.13 (3,778) | 7.61 ± 0.07 (84) | 7.98 ± 0.07 (84) | 0.37 ± 0.07 (84) |
| 7 | Control | 25.7 ± 0.3 (3,736) | 7.98 ± 0.13 (3,771) | 7.80 ± 0.07 (84) | 8.16 ± 0.06 (84) | 0.36 ± 0.06 (84) |
| 8 | OA | 26.0 ± 0.2 (3,452) | 7.78 ± 0.13 (3,839) | 7.61 ± 0.07 (84) | 7.97 ± 0.07 (84) | 0.37 ± 0.07 (84) |

**Table S7** Seawater chemistry in the eight experimental tanks. Values are expressed as mean ± SD with measurement replication (*n*). OA, ocean acidification; $pH_T$, pH on the total scale; TA, total alkalinity; $pCO_2$, partial pressure of $CO_2$; DIC, dissolved inorganic carbon; $\Omega_{ca}$, calcite saturation; $\Omega_{ar}$, aragonite saturation

| Tank | Treatment | Salinity | Temperature (ºC) | $pH_T$ | TA (μmol $kg^{-1}$) | $pCO_2$ (μatm) | DIC (μmol $kg^{-1}$) | $CO_2$ (μmol $kg^{-1}$) | $HCO_3^-$ (μmol $kg^{-1}$) | $CO_3^{2-}$ (μmol $kg^{-1}$) | $\Omega_{ca}$ | $\Omega_{ar}$ |
|---|---|---|---|---|---|---|---|---|---|---|---|---|
| 1 | Control | 34.6 ± 0.4 (10) | 25.9 ± 0.2 (456) | 7.96 ± 0.13 (456) | 2,147 ± 52 (132) | 497 ± 177 (456) | 1,907 ± 75 (456) | 14 ± 5 (456) | 1,715 ± 107 (456) | 179 ± 42 (456) | 4.33 ± 1.01 (456) | 2.86 ± 0.67 (456) |
| 2 | OA | 34.6 ± 0.5 (10) | 25.9 ± 0.2 (366) | 7.77 ± 0.13 (366) | 2,166 ± 54 (120) | 838 ± 292 (366) | 2,018 ± 70 (366) | 23 ± 8 (366) | 1,868 ± 89 (366) | 127 ± 32 (366) | 3.06 ± 0.78 (366) | 2.02 ± 0.52 (366) |
| 3 | Control | 34.6 ± 0.4 (10) | 25.9 ± 0.2 (444) | 7.98 ± 0.12 (444) | 2,166 ± 42 (132) | 472 ± 161 (444) | 1,911 ± 73 (444) | 13 ± 5 (444) | 1,712 ± 105 (444) | 186 ± 41 (444) | 4.50 ± 1.00 (444) | 2.97 ± 0.66 (444) |
| 4 | OA | 34.7 ± 0.4 (10) | 25.9 ± 0.2 (433) | 7.78 ± 0.14 (433) | 2,156 ± 60 (144) | 829 ± 293 (433) | 2,015 ± 75 (433) | 23 ± 8 (433) | 1,864 ± 96 (433) | 128 ± 34 (433) | 3.09 ± 0.82 (433) | 2.04 ± 0.54 (433) |
| 5 | Control | 34.6 ± 0.4 (10) | 26.0 ± 0.3 (341) | 7.99 ± 0.13 (341) | 2,160 ± 54 (120) | 464 ± 167 (341) | 1,902 ± 77 (341) | 13 ± 5 (341) | 1,699 ± 112 (341) | 190 ± 46 (341) | 4.59 ± 1.11 (341) | 3.04 ± 0.74 (341) |
| 6 | OA | 34.7 ± 0.4 (10) | 26.0 ± 0.2 (366) | 7.80 ± 0.13 (366) | 2,146 ± 62 (144) | 779 ± 263 (366) | 1,995 ± 75 (366) | 22 ± 7 (366) | 1,841 ± 95 (366) | 133 ± 34 (366) | 3.21 ± 0.81 (366) | 2.12 ± 0.54 (366) |
| 7 | Control | 34.6 ± 0.4 (10) | 25.7 ± 0.3 (495) | 7.97 ± 0.13 (495) | 2,151 ± 55 (180) | 482 ± 176 (495) | 1,902 ± 80 (495) | 13 ± 5 (495) | 1,707 ± 111 (495) | 182 ± 44 (495) | 4.41 ± 1.06 (495) | 2.91 ± 0.70 (495) |
| 8 | OA | 34.7 ± 0.4 (10) | 25.9 ± 0.2 (338) | 7.79 ± 0.13 (338) | 2,156 ± 60 (120) | 802 ± 292 (338) | 2,006 ± 75 (338) | 22 ± 8 (338) | 1,853 ± 94 (338) | 131 ± 34 (338) | 3.16 ± 0.81 (338) | 2.09 ± 0.54 (338) |

**Table S8** Calcification and surface growth (mean ± SD) of *Acropora cytherea*, *Pocillopora verrucosa*, and *Porites cylindrica* in a control and ocean acidification (OA) treatment. *n*, sample size

| Species | Treatment | Calcification (mg $cm^{-2}$ $day^{-1}$) | Total Calcification (mg $cm^{-2}$) | Surface Growth (% $day^{-1}$) | Total Surface Growth (%) | *n* |
|---|---|---|---|---|---|---|
| *Acropora cytherea* | Control | 0.63 ± 0.29 | 72.93 ± 32.96 | 0.56 ± 0.30 | 70.03 ± 37.39 | 12 |
| *Acropora cytherea* | OA | 0.48 ± 0.25 | 54.40 ± 29.22 | 0.39 ± 0.25 | 49.10 ± 31.97 | 16 |
| *Pocillopora verrucosa* | Control | 2.10 ± 0.53 | 242.05 ± 60.79 | 1.49 ± 0.24 | 187.85 ± 30.26 | 12 |
| *Pocillopora verrucosa* | OA | 1.69 ± 0.42 | 193.79 ± 48.00 | 1.15 ± 0.24 | 144.45 ± 30.25 | 16 |
| *Porites cylindrica* | Control | 1.71 ± 0.61 | 197.80 ± 70.52 | 1.26 ± 0.40 | 157.50 ± 49.39 | 12 |
| *Porites cylindrica* | OA | 1.48 ± 0.49 | 169.40 ± 56.03 | 1.02 ± 0.25 | 127.27 ± 31.28 | 16 |

**Table S9** Post hoc species comparison of linear mixed effects models (LMMs) for calcification and surface growth. Summarised output of LMMs and model information can be found in Table S4. p-values were adjusted with Bonferroni correction. Bold p-values indicate significant effects with $\alpha$ lower than 0.05. Acy, *Acropora cytherea*; Pve, *Pocillopora verrucosa*; Pcy, *Porites cylindrica*

| | Calcification | | | Surface Growth | | |
|---|---|---|---|---|---|---|
| Contrast (Species) | df | t | p | df | t | p |
| Acy vs. Pve | 9.0 | -5.6 | **0.001** | 9.0 | -5.5 | **0.001** |
| Acy vs. Pcy | 9.0 | -4.2 | **0.007** | 9.0 | -4.4 | **0.005** |
| Pve vs. Pcy | 9.0 | 1.4 | 0.577 | 9.0 | 1.2 | 0.819 |

**Table S10** Photosynthesis:Respiration (P:R) ratios of *Acropora cytherea*, *Pocillopora verrucosa*, and *Porites cylindrica* in a control and ocean acidification (OA) treatment measured in low and moderate flow (2 and 6 cm $s^{-1}$, respectively) during the acclimation period ($t_{-1}$, one week before the start of gradual pH decrease) and experimental period ($t_3$, after three weeks under OA conditions, including two weeks of gradual pH decrease; $t_7$, after seven weeks; $t_{12}$, after 12 weeks). Values are means ± SD, with sample size (*n*), and 'pre-OA' refers to the experimental OA group before the start of the OA treatment, with conditions equivalent to the control treatment.

| Species | Time Point | Treatment | Flow | P:R | *n* |
|---|---|---|---|---|---|
| *Acropora cytherea* | $t_{-1}$ | Control | Low | 1.03 ± 0.08 | 12 |
| *Acropora cytherea* | $t_{-1}$ | pre-OA | Low | 1.05 ± 0.15 | 16 |
| *Acropora cytherea* | $t_{-1}$ | Control | Moderate | 0.78 ± 0.14 | 12 |
| *Acropora cytherea* | $t_{-1}$ | pre-OA | Moderate | 1.01 ± 0.15 | 16 |
| *Acropora cytherea* | $t_3$ | Control | Low | 1.26 ± 0.22 | 12 |
| *Acropora cytherea* | $t_3$ | OA | Low | 1.32 ± 0.22 | 16 |
| *Acropora cytherea* | $t_3$ | Control | Moderate | 1.17 ± 0.19 | 12 |
| *Acropora cytherea* | $t_3$ | OA | Moderate | 1.20 ± 0.24 | 16 |
| *Acropora cytherea* | $t_7$ | Control | Low | 1.35 ± 0.17 | 12 |
| *Acropora cytherea* | $t_7$ | OA | Low | 1.26 ± 0.14 | 16 |
| *Acropora cytherea* | $t_7$ | Control | Moderate | 1.43 ± 0.25 | 12 |
| *Acropora cytherea* | $t_7$ | OA | Moderate | 1.41 ± 0.21 | 16 |
| *Acropora cytherea* | $t_{12}$ | Control | Low | 1.26 ± 0.09 | 12 |
| *Acropora cytherea* | $t_{12}$ | OA | Low | 1.06 ± 0.11 | 16 |
| *Acropora cytherea* | $t_{12}$ | Control | Moderate | 1.51 ± 0.20 | 12 |
| *Acropora cytherea* | $t_{12}$ | OA | Moderate | 1.16 ± 0.10 | 16 |
| *Pocillopora verrucosa* | $t_{-1}$ | Control | Low | 1.27 ± 0.15 | 12 |
| *Pocillopora verrucosa* | $t_{-1}$ | pre-OA | Low | 1.54 ± 0.22 | 16 |
| *Pocillopora verrucosa* | $t_{-1}$ | Control | Moderate | 1.08 ± 0.09 | 12 |
| *Pocillopora verrucosa* | $t_{-1}$ | pre-OA | Moderate | 1.14 ± 0.14 | 16 |
| *Pocillopora verrucosa* | $t_3$ | Control | Low | 1.58 ± 0.15 | 12 |
| *Pocillopora verrucosa* | $t_3$ | OA | Low | 1.61 ± 0.16 | 16 |
| *Pocillopora verrucosa* | $t_3$ | Control | Moderate | 1.17 ± 0.21 | 12 |
| *Pocillopora verrucosa* | $t_3$ | OA | Moderate | 1.09 ± 0.16 | 16 |
| *Pocillopora verrucosa* | $t_7$ | Control | Low | 1.42 ± 0.13 | 12 |
| *Pocillopora verrucosa* | $t_7$ | OA | Low | 1.34 ± 0.14 | 16 |
| *Pocillopora verrucosa* | $t_7$ | Control | Moderate | 1.30 ± 0.12 | 12 |
| *Pocillopora verrucosa* | $t_7$ | OA | Moderate | 1.26 ± 0.13 | 16 |
| *Pocillopora verrucosa* | $t_{12}$ | Control | Low | 1.38 ± 0.23 | 12 |
| *Pocillopora verrucosa* | $t_{12}$ | OA | Low | 1.29 ± 0.17 | 16 |
| *Pocillopora verrucosa* | $t_{12}$ | Control | Moderate | 1.27 ± 0.15 | 12 |
| *Pocillopora verrucosa* | $t_{12}$ | OA | Moderate | 1.11 ± 0.14 | 16 |
| *Porites cylindrica* | $t_{-1}$ | Control | Low | 0.96 ± 0.23 | 12 |
| *Porites cylindrica* | $t_{-1}$ | pre-OA | Low | 1.28 ± 0.19 | 16 |
| *Porites cylindrica* | $t_{-1}$ | Control | Moderate | 1.14 ± 0.24 | 12 |
| *Porites cylindrica* | $t_{-1}$ | pre-OA | Moderate | 1.21 ± 0.37 | 16 |
| *Porites cylindrica* | $t_3$ | Control | Low | 0.98 ± 0.19 | 12 |
| *Porites cylindrica* | $t_3$ | OA | Low | 1.11 ± 0.17 | 16 |
| *Porites cylindrica* | $t_3$ | Control | Moderate | 1.46 ± 0.19 | 12 |

| | | | | | |
|---|---|---|---|---|---|
| *Porites cylindrica* | $t_3$ | OA | Moderate | 1.13 ± 0.23 | 16 |
| *Porites cylindrica* | $t_7$ | Control | Low | 1.32 ± 0.10 | 12 |
| *Porites cylindrica* | $t_7$ | OA | Low | 1.40 ± 0.14 | 16 |
| *Porites cylindrica* | $t_7$ | Control | Moderate | 1.63 ± 0.18 | 12 |
| *Porites cylindrica* | $t_7$ | OA | Moderate | 1.41 ± 0.15 | 16 |
| *Porites cylindrica* | $t_{12}$ | Control | Low | 1.12 ± 0.18 | 12 |
| *Porites cylindrica* | $t_{12}$ | OA | Low | 1.00 ± 0.21 | 16 |
| *Porites cylindrica* | $t_{12}$ | Control | Moderate | 1.29 ± 0.27 | 12 |
| *Porites cylindrica* | $t_{12}$ | OA | Moderate | 0.92 ± 0.21 | 16 |

**Table S11** Post hoc analyses of species' linear mixed effects models (LMMs) for photosynthesis:respiration (P:R) ratio. Summarised output of LMMs and model information can be found in Table S5. p-values were adjusted with Bonferroni correction. Bold p-values indicate significant effects with $\alpha$ lower than 0.05. $t_3$, after three weeks under ocean acidification (OA) conditions, including two weeks of gradual pH decrease; $t_7$, after seven weeks; $t_{12}$, after 12 weeks; LF, low flow; MF, moderate flow

| | | *Acropora cytherea* | | | *Pocillopora verrucosa* | | | *Porites cylindrica* | | |
|---|---|---|---|---|---|---|---|---|---|---|
| | | df | t | p | df | t | p | df | t | p |
| Contrast (Time) | | | | | | | | | | |
| | $t_3$ vs. $t_7$ | 130.0 | -4.0 | **< 0.001** | 130.0 | 1.3 | 0.625 | 130.0 | -7.8 | < 0.001 |
| | $t_3$ vs. $t_{12}$ | 130.0 | -0.3 | 1.000 | 130.0 | 4.3 | **< 0.001** | 130.0 | 2.5 | **0.037** |
| | $t_7$ vs. $t_{12}$ | 130.0 | 3.7 | **0.001** | 130.0 | 3.0 | **0.010** | 130.0 | 10.4 | **< 0.001** |
| Contrast (Treatment:Flow) | | | | | | | | | | |
| | Control LF vs. OA LF | 6.4 | 1.1 | 0.645 | 5.9 | 0.6 | 1.000 | 9.3 | -0.6 | 1.000 |
| | Control MF vs. OA MF | 6.4 | 1.5 | 0.373 | 5.9 | 1.4 | 0.449 | 9.3 | 5.6 | **0.001** |
| Contrast (Treatment:Time) | | | | | | | | | | |
| | Control $t_3$ vs. OA $t_3$ | 8.0 | -0.5 | 1.000 | 6.9 | 0.3 | 1.000 | 14.7 | 1.7 | 0.360 |
| | Control $t_7$ vs. OA $t_7$ | 8.0 | 0.6 | 1.000 | 6.9 | 0.8 | 1.000 | 14.7 | 1.2 | 0.787 |
| | Control $t_{12}$ vs. OA $t_{12}$ | 8.0 | 3.5 | **0.024** | 6.9 | 1.8 | 0.357 | 14.7 | 3.9 | **0.004** |
| Contrast (Flow:Time) | | | | | | | | | | |
| | LF $t_3$ vs. MF $t_3$ | 130.0 | 2.3 | 0.062 | 130.0 | 13.6 | **< 0.001** | 130.0 | -5.1 | **< 0.001** |
| | LF $t_7$ vs. MF $t_7$ | 130.0 | -2.6 | **0.032** | 130.0 | 2.9 | **0.013** | 130.0 | -3.3 | **0.004** |
| | LF $t_{12}$ vs. MF $t_{12}$ | 130.0 | -4.1 | **< 0.001** | 130.0 | 4.3 | **< 0.001** | 130.0 | -1.0 | 0.960 |
| Contrast (Treatment:Flow:Time) | | | | | | | | | | |
| | Control LF $t_3$ vs. OA LF $t_3$ | 13.6 | -0.6 | 1.000 | 10.2 | -0.5 | 1.000 | 35.5 | -1.6 | 0.680 |
| | Control MF $t_3$ vs. OA MF $t_3$ | 13.6 | -0.4 | 1.000 | 10.2 | 1.1 | 1.000 | 35.5 | 4.2 | **0.001** |
| | Control LF $t_7$ vs. OA LF $t_7$ | 13.6 | 1.0 | 1.000 | 10.2 | 1.0 | 1.000 | 35.5 | -1.1 | 1.000 |
| | Control MF $t_7$ vs. OA MF $t_7$ | 13.6 | 0.1 | 1.000 | 10.2 | 0.5 | 1.000 | 35.5 | 2.9 | **0.039** |
| | Control LF $t_{12}$ vs. OA LF $t_{12}$ | 13.6 | 2.2 | 0.251 | 10.2 | 1.2 | 1.000 | 35.5 | 1.5 | 0.903 |
| | Control MF $t_{12}$ vs. OA MF $t_{12}$ | 13.6 | 3.9 | **0.010** | 10.2 | 2.0 | 0.418 | 35.5 | 4.7 | **< 0.001** |

### Supplementary Figures

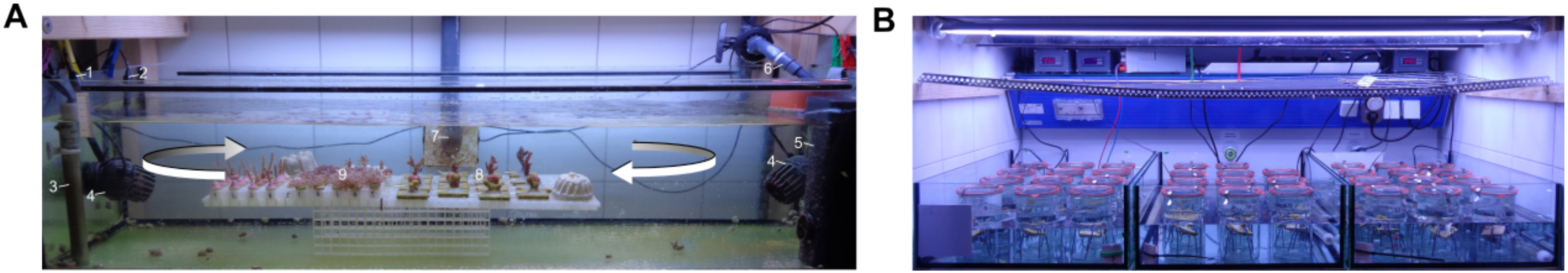

**Fig. S1** Experimental setup. (A) Detail of an experimental tank with the main elements marked: (1) pH probe, (2) temperature probe, (3) heater, (4) pump, (5) wave generator, (6) inflow, (7) outflow, (8) scleractinian corals: (front to back) *Porites cylindrica, Pocillopora verrucosa, Acropora cytherea, Montipora digitata*, (9) octocorals: (front to back) *Xenia umbellata*, *Pinnigorgia flava*, *Sinularia* sp., *Plexaurella* sp. Arrows indicate the direction of water circulation. (B) Overview of the incubation setup of respirometry assays.

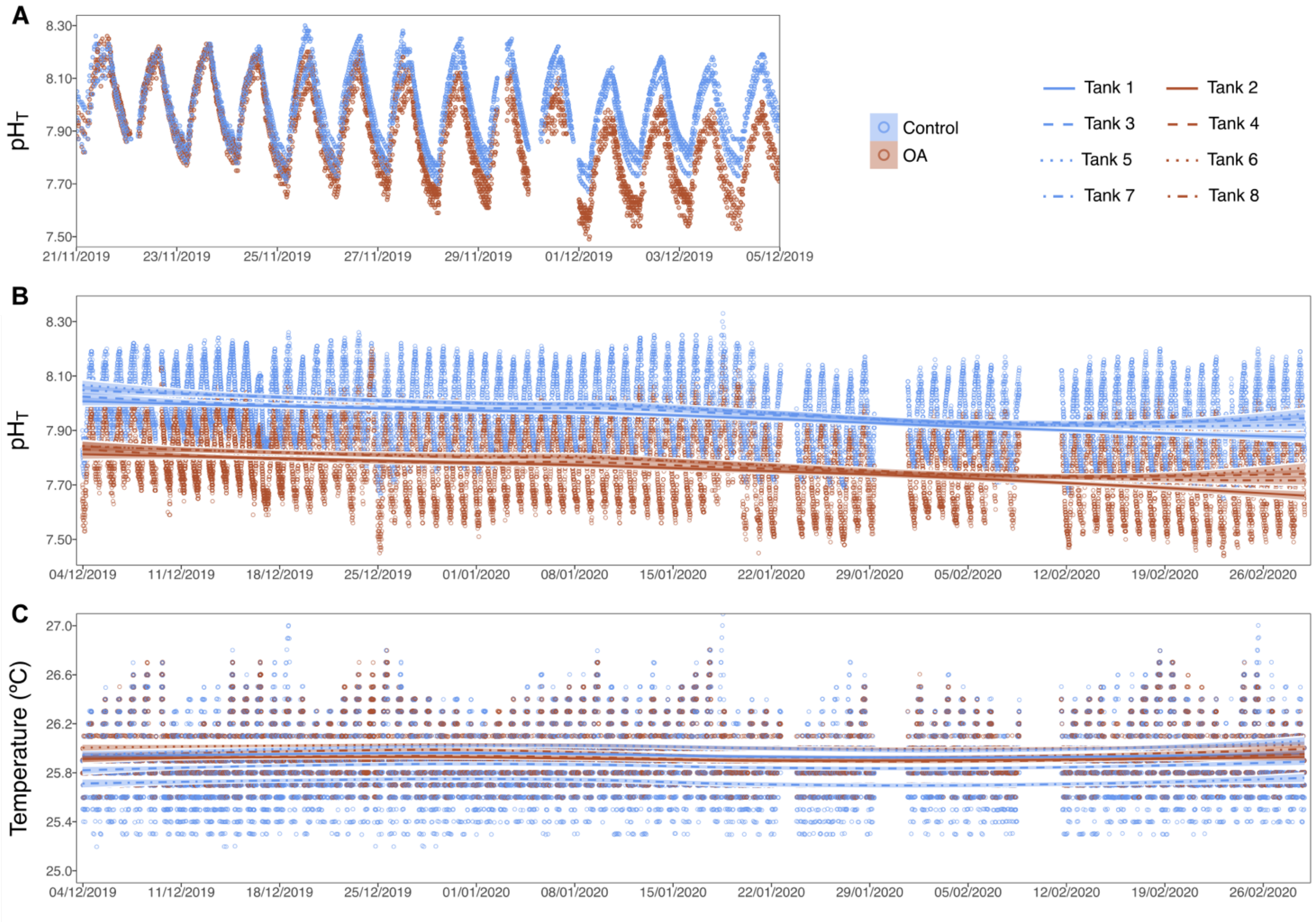

**Fig. S2** Seawater conditions during the experiment. (A) pH during gradual establishment of ocean acidification (OA) treatment. (B) pH and (C) temperature during the experiment after reaching target OA conditions in OA treatment tanks. Lines are local polynomial regression fitting with 95% confidence intervals (shaded area). Dates are provided as DD/MM/YYYY. Data gaps reflect periods when values were not stored, which did not affect the measurement and manipulation of pH and temperature in the tanks, except on 09/12/2019 when the digital controller of the experimental system experienced a power failure. This stoppage lasted < 9 h, during which pH in the OA treatment tanks reverted to ambient levels and upon restarting the system, pH was gradually restored to target levels. pH$_T$, pH on the total scale

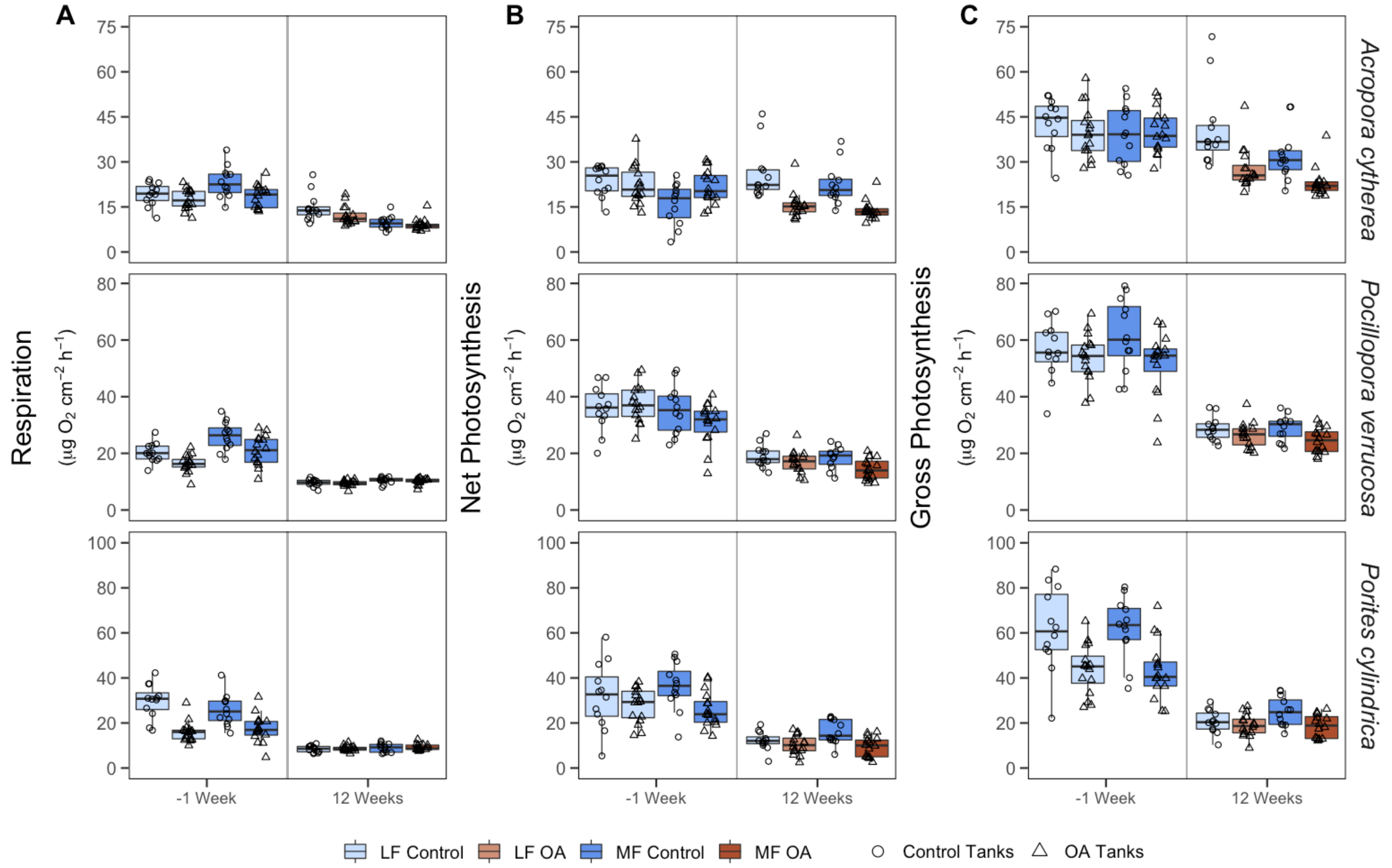

**Fig. S3** Physiological responses in respiration and photosynthesis of three reef-building coral species. Rates of (A) respiration, (B) net photosynthesis, and (C) gross photosynthesis of *Acropora cytherea*, *Pocillopora verrucosa*, and *Porites cylindrica* in a control and ocean acidification (OA) treatment measured in low flow (LF, 2 cm $s^{-1}$) and moderate flow (MF, 6 cm $s^{-1}$) conditions during the acclimation period (measured one week before the start of the gradual pH decrease) and after 12 weeks under OA conditions, including two weeks of gradual pH decrease. Point shapes indicate treatment assignment. Data from the acclimation period are presented separated by the respective treatment applied during the OA phase. Boxes represent the first and third quartiles with lines as medians and whiskers as the minimum and maximum values or up to the 1.5 × interquartile range (IQR), whichever is reached first.

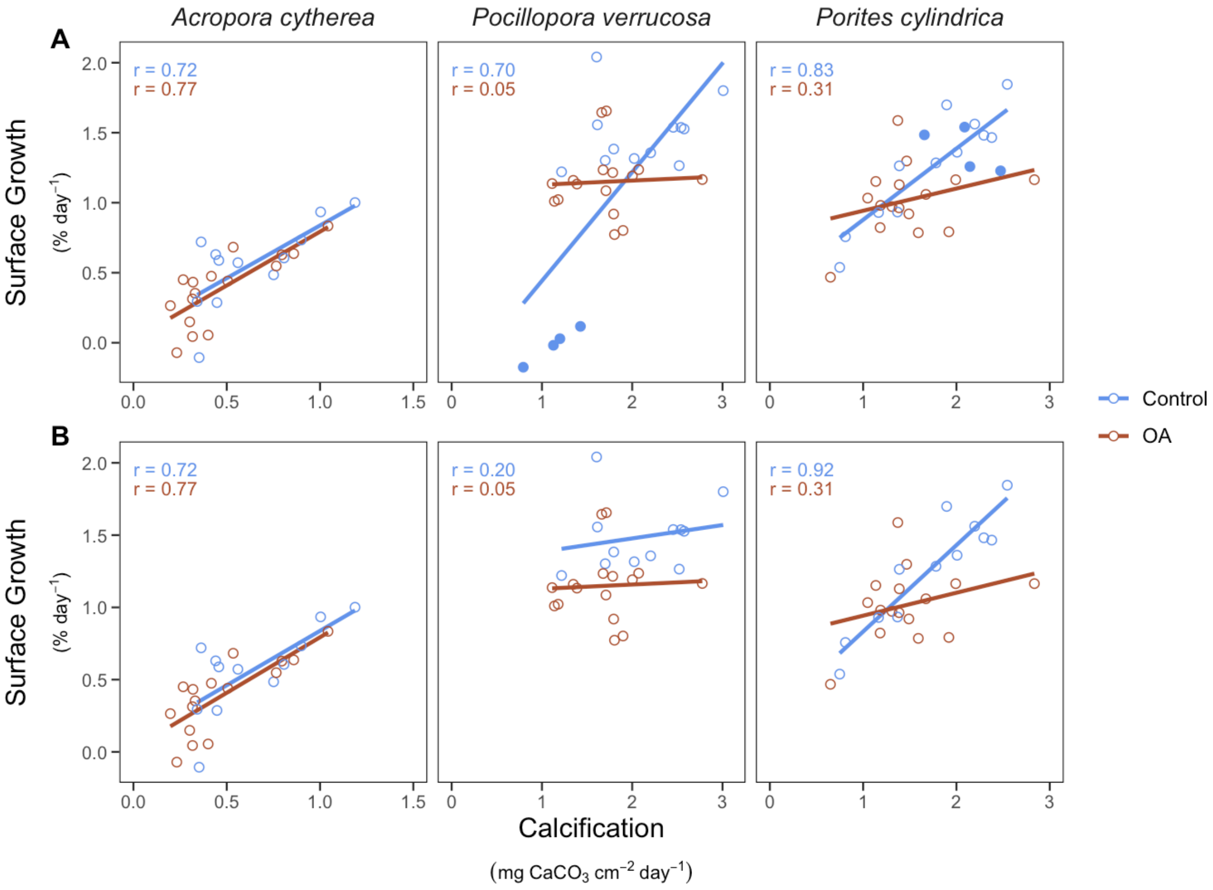

**Fig. S4** Scatterplot of surface growth against calcification of *Acropora cytherea*, *Pocillopora verrucosa*, and *Porites cylindrica* after three months in a control and ocean acidification (OA) treatment. (A) Complete dataset. (B) Dataset with excluded fragments from the affected experimental tank (i.e., tank five) (see Materials and Methods). Coloured points indicate values from experimental tank five. Lines are linear regression fitting. r, Pearson correlation coefficient
